# Blood immunophenotyping identifies distinct kidney histopathology and outcomes in patients with lupus nephritis

**DOI:** 10.1101/2024.01.14.575609

**Authors:** Alice Horisberger, Alec Griffith, Joshua Keegan, Arnon Arazi, John Pulford, Ekaterina Murzin, Kaitlyn Howard, Brandon Hancock, Andrea Fava, Takanori Sasaki, Tusharkanti Ghosh, Jun Inamo, Rebecca Beuschel, Ye Cao, Katie Preisinger, Maria Gutierrez-Arcelus, Thomas M. Eisenhaure, Joel Guthridge, Paul J. Hoover, Maria Dall’Era, David Wofsy, Diane L. Kamen, Kenneth C. Kalunian, Richard Furie, Michael Belmont, Peter Izmirly, Robert Clancy, David Hildeman, E. Steve Woodle, William Apruzzese, Maureen A. McMahon, Jennifer Grossman, Jennifer L. Barnas, Fernanda Payan-Schober, Mariko Ishimori, Michael Weisman, Matthias Kretzler, Celine C. Berthier, Jeffrey B. Hodgin, Dawit S. Demeke, Chaim Putterman, Accelerating Medicines Partnership: RA/SLE Network, Michael B. Brenner, Jennifer H. Anolik, Soumya Raychaudhuri, Nir Hacohen, Judith A. James, Anne Davidson, Michelle A. Petri, Jill P. Buyon, Betty Diamond, Fan Zhang, James A. Lederer, Deepak A. Rao

## Abstract

Lupus nephritis (LN) is a frequent manifestation of systemic lupus erythematosus, and fewer than half of patients achieve complete renal response with standard immunosuppressants. Identifying non-invasive, blood-based pathologic immune alterations associated with renal injury could aid therapeutic decisions. Here, we used mass cytometry immunophenotyping of peripheral blood mononuclear cells in 145 patients with biopsy-proven LN and 40 healthy controls to evaluate the heterogeneity of immune activation in patients with LN and to identify correlates of renal parameters and treatment response. Unbiased analysis identified 3 immunologically distinct groups of patients with LN that were associated with different patterns of histopathology, renal cell infiltrates, urine proteomic profiles, and treatment response at one year. Patients with enriched circulating granzyme B^+^ T cells at baseline showed more severe disease and increased numbers of activated CD8 T cells in the kidney, yet they had the highest likelihood of treatment response. A second group characterized primarily by a high type I interferon signature had a lower likelihood of response to therapy, while a third group appeared immunologically inactive by immunophenotyping at enrollment but with chronic renal injuries. Main immune profiles could be distilled down to 5 simple cytometric parameters that recapitulate several of the associations, highlighting the potential for blood immune profiling to translate to clinically useful non-invasive metrics to assess immune-mediated disease in LN.

## Introduction

Lupus nephritis (LN) is a common and severe manifestation of systemic lupus erythematosus (SLE) that affects over 40% of patients(1) and substantially increases SLE-associated morbidity and mortality(2–4). Advances in the understanding of renal inflammatory processes in patients with severe LN(5–9) have not yet translated into improved renal outcomes(10). Less than half of the patients treated with standard immunosuppressive regimens achieve complete renal response at one year, and 10% to 30% progress to end-stage renal disease(10–12). The heterogeneity of immune pathways involved in patients with SLE and LN contributes to the variability in treatment response and poses challenges in developing successful new therapies(13, 14).

Current stratification of patients with LN relies on histologic classification of the nature, intensity, and localization of glomerular involvement (ISN/RPS class), as well as the degree of active inflammation and fibrosis in the glomeruli and tubular-interstitium (NIH activity and chronicity indices)(15). These features currently guide treatment decisions(16, 17) and inclusion criteria in clinical trials (class III, IV and/or V). However they have several limitations: they do not inform on the inflammatory pathways involved, they have limited ability to predict long-term renal outcome, and they show discordance with clinical response(7, 18). Stratifying patients based on molecular or cellular features of immune-mediated pathways would improve the predictive value of kidney biopsy(19). Efforts to characterize inflammatory pathways involved in the kidney tissue in lupus nephritis have identified signatures associated with poor clinical outcome, such as accumulation of CD4^neg^ T cells(20), enriched fibrotic pathways, type I interferon signaling(6), and specific proinflammatory cytokines and chemokines(7, 21).

In parallel with analyses of kidney biopsy tissue, identifying non-invasive, easily accessible methods to characterize LN immune heterogeneity and track LN activity could substantially improve disease management. Prior efforts to characterize immunologic diversity among patients with SLE, not restricted to LN, using blood transcriptomics and proteomics have identified distinct signatures. Evidence of upregulated type I Interferon (IFN) signalling as measured by IFN response gene signatures has been repeatedly identified in 50-80% of patients with SLE and is associated with more severe disease, including renal manifestations(22, 23). Independently, a circulating neutrophil signature has been associated with active renal involvement in patients with SLE(24, 25) and appears to improve with renal response to treatment(26, 27). More generally, SLE disease activity has been characterized by specific transcriptomic signatures involving myeloid, plasmablast(24, 25, 28), B-cell activation, and cell cycle pathways(25). Similarly, cytometric analyses have highlighted expanded activated B cells, plasmablasts, B-helper T cells and activated myeloid cells in the blood of patients with active SLE, including patients with LN(29–33). Clustering patients with SLE based on blood immunophenotyping by flow cytometry has shown promise in identifying distinct groups of patients with SLE with different treatment responses(34). However, the extent to which these cellular signatures can inform renal pathology and treatment response across patients with LN is unclear.

Accordingly, this study was initiated to identify patterns of immune cell signatures in the blood of patients with LN that are associated with renal tissue activity and treatment response. By applying mass cytometry immunophenotyping to a prospective cohort of patients with biopsy proven LN in a multi-ethnic/racial study across the U.S., we have defined alterations in circulating T cells, B cells, myeloid cells and NK cells. Using classifications and unsupervised models, cellular profiles in blood have been identified that discriminate three groups of patients differing in renal activity and outcome. Further using paired single cell RNA-seq of kidney immune cells and proteomic analyses of urine, we found that these blood immunophenotype-defined groups were associated with specific cell infiltrates in kidneys and proteomic signatures in the urine.

## Results

### Overview of blood immunophenotyping in a longitudinal cohort of patients with active LN

We analyzed peripheral blood mononuclear cells (PBMCs) by mass cytometry using four 48-marker panels in 145 patients with SLE and biopsy-proven LN class III, IV, and/or V, as well as 40 healthy controls enrolled in the AMP SLE phase II study (**Supplemental Tables 1 and 2**). Compared to the controls, patients with LN in this study were younger (p=0.02) and more frequently female, Black (p=0.003), and Hispanic or Latino (p=0.008). Within patients with LN, 68% had positive anti-dsDNA antibody and/or low serum complement levels, and 50% had extrarenal clinical manifestations of SLE at the time of biopsy. All patients had a clinical indication for renal biopsy based on proteinuria (defined as urine protein to creatinine ratio (UPCR) > 0.5 g/g)(16, 17). Most biopsies reported a proliferative classification (III or IV +/- V, 70%) compared to pure membranous (V, 30%). Sixty-nine percent of the patients had a history of previous renal biopsy; 61% had one or more prior episodes of biopsy-proven proliferative or membranous LN. Patients were treated following standard of care according to the judgment of their physician(16, 17). Clinical renal response was defined as previously reported in patients with a UPCR ≥ 1 at baseline(35, 36). At 52 weeks, 28% achieved complete response, 24% a partial response and 48% no response. Longitudinal blood samples from 50 patients were analyzed at week-12 and/or week-52 to determine the trajectory of immune cells following treatment.

To minimize technical confounders, PBMC samples were cryopreserved with optimized protocol(37) and then processed for mass cytometry at a single central site; LN and control samples were balanced across 23 batches. Each PBMC sample was stained with panels designed to interrogate T, B, myeloid, or NK cells; a total of 267 samples were retained after quality control (**Supplemental Table 2, Supplemental Figure 1A**). A total of 25-30×10^6^ cells/panel were available for analysis after quality control, averaging approximately 111,000 cells/sample/panel (**Supplemental Figure 1B**). Batch effect correction with Harmony(38) reduced inter-batch signal variation as measured by the local inverse Simpson’s index, without impairing cell type identification or removing signal variation related to LN versus control (**Supplemental Figures 1C-E**). We then applied a stepwise unsupervised approach to analyze the data generated from each panel, starting from total PBMCs, which avoids potential bias from manual gating. To ensure equal representation of each sample, we downsampled randomly to 10,000 cells/sample. We labeled each cluster as either T, B, myeloid or NK cells, based on cell-type canonical markers, and projected the results in a uniform manifold approximation and projection (UMAP) (**Figure 1A, Supplemental Figure 1F**). We then extracted the major cell lineage of interest (T, B, myeloid, and NK) from the dedicated panel and re-clustered the cells to define fine-grained cell-type specific neighborhoods and clusters (**Figure 1B, Supplemental Figure 2**).

**Figure 1.**
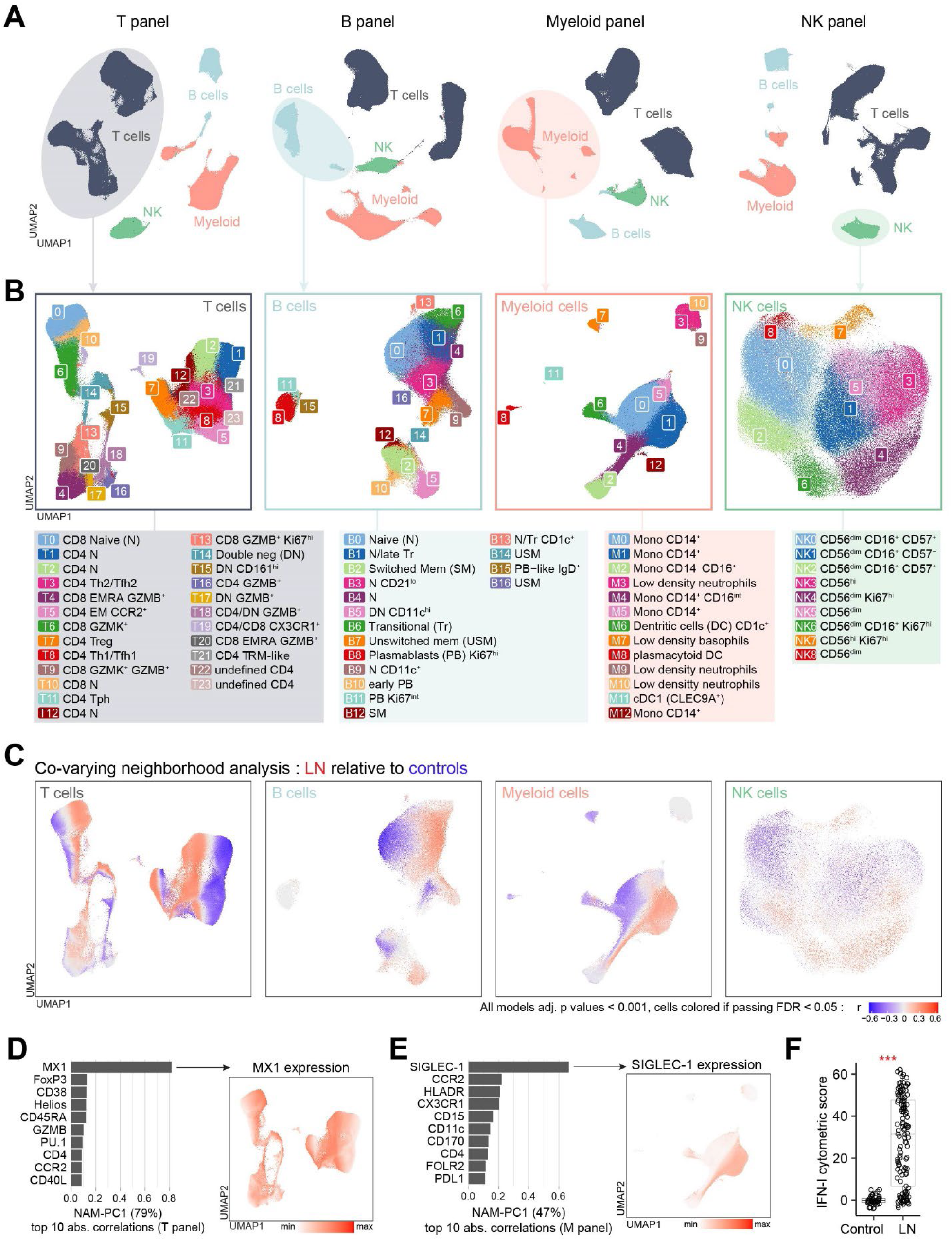
Blood immunophenotyping by mass cytometry captures the range of type I interferon (IFN) signaling intensity in lupus nephritis (LN). (**A**) Identification of major immune cell types within peripheral blood mononuclear cells from 267 samples (from 145 patients with LN and 40 controls) stained with four different panels. (**B**) Detailed cell type specific clustering analysis in each dedicated panel (e.g. T cells in the T panel). (**C**) Cell-type specific associations with patients with LN relative to controls at baseline, using covarying neighborhood analysis (CNA) and adjusted for demographic variables (age, sex, ethnicity and race). Cells in the UMAP are expanded (red) or depleted (blue) in patients with LN. (**D**) Top 10 correlation coefficients between markers in the T panel or (**E**), the Myeloid panel and the major axis of variation of CNA, defined by the first principal component analysis (PC1) on a matrix including all neighborhood abundances (NAM). The correlation coefficient was defined using Spearman’s rho testing. (**F**), Comparison of a type I IFN cytometric score between LN (n=125) and controls (n=40). Statistical significance was determined using Mann-Whitney U test, ***p<0.001.

To validate the robustness of our unbiased analysis, we confirmed our ability to reproduce previously reported alterations in immune cell subset frequencies in patients with SLE. We identified alterations in the frequency of rare cell populations including plasmablasts (CD20^-^ CD19^int^CD27^hi^CD38^hi^)(32) and plasmacytoid dendritic cells (pDC, CD123^+^HLADR^+^IRF8^+^)(39, 40), with abundances similar to prior reports (**Supplemental Figure 1G-H**). We also confirmed in our cohort an expansion of cell subsets previously associated with LN, including T peripheral helper cells (Tph, cluster T11, CD4^+^PD1^+^ICOS^+^CXCR5^-^)(32) and CD11c^+^ B cell subsets (clusters B5 and B9, CD11c^+^T-bet^+^CD21^-^IgD^+/-^)(29, 30, 33), independently from demographic variables (age, sex, ethnicity and race) and immunosuppressive therapy (**Supplemental Figure 1I**).

### A cytometric score captures type I interferon signature heterogeneity in LN

To explore blood immune cell alterations in patients with LN comprehensively, we applied a covarying neighborhood analysis (CNA) in each panel to identify disease-associated cells in a cluster-free manner, while adjusting for sample-level covariates(41). CNA results were consistent with our initial analysis of Tph and activated B cells, yet additionally detected marked diffuse alterations within each cell type, after adjusting for demographic variables and accounting for false discovery rate (FDR) < 0.05, suggesting broad differences in the baseline cytometric profiles of patients with LN compared to controls (**Figure 1C**). CNA results strongly correlated with those from a cluster-based single-cell mixed-effect model (MASC)(42), providing higher granularity of cell-level association, and were not affected by small cluster outliers (**Supplemental Figures 3A-B**). Patterns of LN-associated cells were maintained when comparing patients not on immunosuppressive/biologic treatment and on <= 5mg of prednisone at baseline with controls, with striking differences particularly in the T cells and myeloid cells, despite reduced sample size (**Supplemental Figure 3C**).

We then examined which markers explained the differences observed between LN and controls. Using the results generated from CNA analysis, we were able to identify the proteins associated with the axes of greatest variation characterized in the NAM (cell neighborhood abundances)-derived principal components. In the T panel, the first principal component (NAM-PC1) explained the majority (79%) of the variance observed and was conspicuously correlated with MX1, a type I interferon (IFN)-induced protein(43), compared to the other markers (absolute r=0.82, p<0.001) (**Figure 1D**). In parallel, in the myeloid panel, monocyte expression of SIGLEC-1, an IFN-inducible protein, was strongly associated with CNA defined LN-enriched cells (**Figure 1E**). Moreover, in the B cell panel, ISG-15, a third type I IFN-induced marker(24, 25, 44), was also increased in all cell types from patients with LN and its expression strongly correlated with MX1 (r=0.89, p<0.001) and SIGLEC-1 (r=0.75, p<0.001) at a sample level (**Supplemental Figure 3D**). MX1 and ISG15 were enriched in all cell types from patients with LN compared to controls, whereas SIGLEC-1^hi^ cells were only detected in myeloid cells, consistent with previous transcriptomic studies(25, 45) (**Supplemental Figure 3E**).

Given the importance of type I IFN signaling in SLE and the emergence of therapeutic IFN receptor blockade(19, 46, 47), we sought to define a cytometric IFN score based on the expression of these three proteins (MX1, SIGLEC-1, and ISG-15, see **Method**). We confirmed a significant increase in cytometric IFN score in patients with LN at baseline compared to controls (**Figure 1F**). In addition, the type I IFN signaling was associated with the presence of serologic manifestations (presence of anti-dsDNA and low complement, **Supplemental Figure 3F**), consistent with results from gene scores in SLE cohorts(47, 48) and with the mechanistic connection between immune complexes inducing type I IFN release, further promoting auto-antibody production(49). While type I interferon activation pathway is associated with SLE severity, including LN, its correlation with renal histology is less clear(6, 23, 47, 50). In this LN cohort, the cytometrically-derived type I IFN score did not correlate with proliferative class, renal activity by NIH index, including the interstitial component, or extrarenal clinical manifestations (**Supplemental Figures 3G-H**). IFN score was also not associated with baseline prednisone dose or the use of immunosuppressants (**Supplemental Figure 3G**). These results indicate that type I IFN signaling was common across clinically and histologically heterogeneous patients with LN.

### Circulating immune cell alterations are associated with histologic patterns of active LN

As the use of immunosuppressive treatment in patients with LN is currently driven by the biopsy-defined proliferative class and the extent of renal activity(16, 17, 51), we next set out to investigate whether immune alterations in the blood reflected biopsy class and features of active renal inflammation. Both proliferative LN classes (III or IV +/- V) and renal activity were predominantly associated with shifts in naïve B cell phenotypes, after adjusting for demographic features and the history of previous renal biopsy (**Figures 2A-B**, **Supplemental Figures 4A-B**). Furthermore, differences in circulating naive B cells were associated with specific histologic features of the activity index, including glomerular endocapillary hypercellularity and the presence and degree of fibrinoid necrosis. The association between naive B cells and fibrinoid necrosis was still evident even when evaluating only the 15 patients with LN who had no immunosuppressant treatment and prednisone ≤ 5mg at the time of sampling (**Supplemental Figures 4C-D**). We also detected a strong association between cells with a neutrophil phenotype (M3, CD14^-^CD16^+^CD11b^+^CD15^hi^), likely representing low-density neutrophils(52), and fibrinoid necrosis, even when including only non-immunosuppressed (with maximum 5mg of prednisone) patients (**Figure 2C, Supplemental Figures 4C-D**). When examining only non-immunosuppressed patients, NIH activity index and fibrinoid necrosis were also associated with a subset of CD14^+^monocytes characterized by low expression of CD86 (M5), a protein downregulated with LPS stimulation and sepsis(53, 54), suggesting a inflammatory state. While B and myeloid cells showed an association with glomerular injuries, shifts within the T cells correlated with the interstitial component of the NIH activity index (**Figure 2D**).

**Figure 2.**
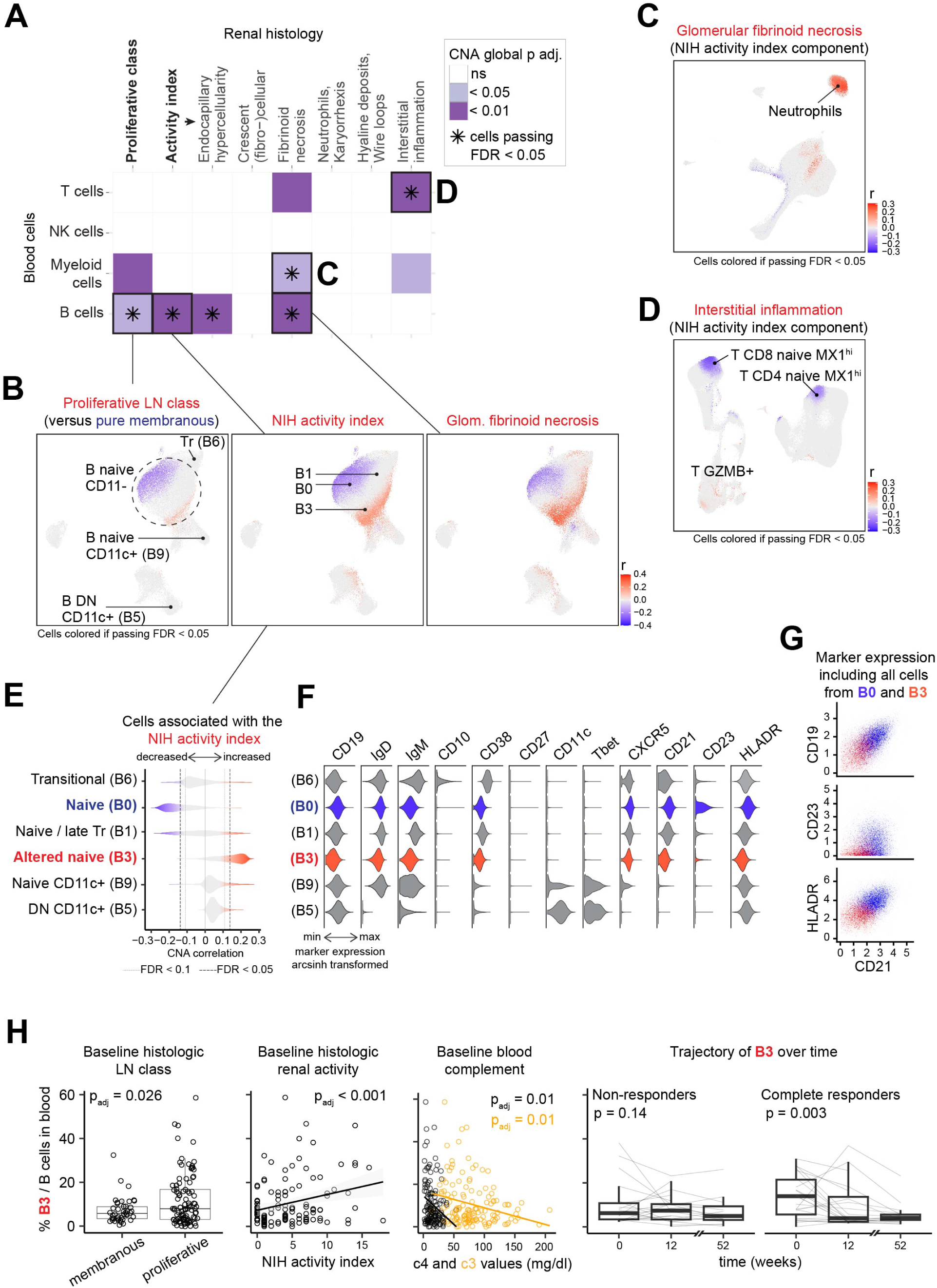
Circulating immune cell subsets association with histologic patterns of active lupus nephritis (LN). (**A**) Summary of the results testing the association within each blood cell type (y axis) and different histologic patterns of LN disease (x axis). Statistical significance was determined using covarying neighborhood analysis (CNA) after adjusting for demographic variables (age, sex, ethnicity and race) and history of previous biopsy. The color represent the global CNA p value as indicated and the star indicates if local associations passed FDR < 0.05. (**B**) Selection of detailed CNA results between B cell alterations, (**C**) myeloid alterations, and (**D**) T cell alterations and specified histologic characteristics. (**E**) Distribution of NIH activity index-associated cell neighborhood correlations (detailed CNA results) in a selection of B cell clusters. (**F**) Violin plots of selected proteins expression in selected B cell clusters. (**G**) Scatterplot of cluster B0 and B3 expression of selected proteins. (**H**) Proportion of cluster B3 (among B cells) association with LN histologic class (n=140), activity index (n=124) and complement levels (n=138), as well as longitudinal change in non-responders (n=19) and complete responders (n=23). Statistical significance was determined using linear models adjusting for demographic variables and history of previous biopsy for cross-sectional analysis at baseline, and using mixed effect models for longitudinal analysis to include patients as a random effect.

Among B cells, the positive association with different renal inflammatory lesions was primarily driven by cluster B3, a naive subset (IgD^+^IgM^+^CD27^-^) with features distinct from transitional cells (B6) and from activated CD11c^+^ subsets (B5, B9), including lack of CD11c, T-bet, CD10, and intermediate CD38 expression (**Figures 2E-F**). Cluster B3 lacked CD23 expression and had a lower expression of CD21 and CD19 compared to the predominant naïve B cell cluster observed in controls (B0) (**Figures 2F-G**). Despite the absence of activation or proliferation markers (HLA-DR, CD95, CD86 or Ki67), this cluster B3 was significantly associated with the NIH activity index and low complement levels (e.g., C3 and C4), independently from demographic variables and history of previous biopsies (**Figure 2H**). In contrast, B3 was not associated with extrarenal manifestations of disease activity (presence of clinical extrarenal SLEDAI, p=0.36). Longitudinal analysis showed a decrease in the proportion of cluster B3 over time in the patients with a complete renal response at 52 weeks, but not in patients with no response (**Figure 2H**). By manual gating, decreased expression of CD21 within naïve B cells (median CD21 within CD20^+^CD27^-^IgD^+^ CD38^int/-^CD10^-^ CD11c^-^ cells) confirmed: 1) an association with active and proliferative LN, 2) an association with low complement levels, and 3) an increased CD21 expression over time in complete responders (**Supplemental Figures 5A-B**). Together, these results support a shift in naïve B cells towards an altered phenotype in patients with LN with active renal inflammation and low complement levels.

### Cytotoxic T cells, proliferating T-B subsets and type I interferon characterize diverse cellular signatures in patients with LN

These results highlight relationships between blood immune cells and histologic evidence of active renal injury, yet the limited ability of histologic classification to predict renal outcome, including in the AMP phase II cohort(36), and the immune heterogeneity observed in the cohort challenged us to approach the data in an unsupervised manner to identify cellular profiles associated with clinical features of LN. As a first step, we presented the global relationships between immune cell subsets in patients with LN. By applying clustering to the correlation matrix of the identified immune cell clusters (after excluding small cluster outliers, see **Method**) and type I IFN score from LN baseline samples (n=115 stained with four panels), we observed sets of coordinated immune cell abundances (**Figure 3A**, **Supplemental Figures 6A**). As expected, cell subsets marked by type I IFN-induced proteins, such as SIGLEC-1^hi^ monocytes (M1) and MX1^hi^ naive CD4 T cells (T2 and T12), were part of a core set correlating with the cytometric type I IFN score (**Figure 3A**). Interestingly, T12 was phenotypically distinct from T2 by its reduced expression of TCF1, a key transcription factor in T cell stemness that is downregulated upon CD4^+^ effector T cell differentiation(55, 56) (**Supplemental Figures 6B-C**). Supporting the hypothesis that these cells were recently activated in an environment enriched in type I IFN, we used a published bulk RNA-seq dataset(57) and found that anti-CD3/CD28 stimulation of naive CD4 T cells downregulated *TCF7* gene (encoding TCF1) expression, and this was accelerated by IFN-β (**Supplemental Figure 6D**). Other cells correlating and closely organized with the type I IFN core set included MX1^hi^ TCF1^lo^ naive CD8 T cells (T10), Ki67^hi^ NK cell subsets (NK4, NK6, NK7), transitional B cells (B6), and a small cluster of B cells with features of marginal zone precursors(58) (B13, IgM^+^CD38^+^CD27^-^CD1c^+^), consistent with the previously reported roles of type I IFN in NK cell activation(59) and in immature B cell expansion(60) (**Figure 3A**).

**Figure 3.**
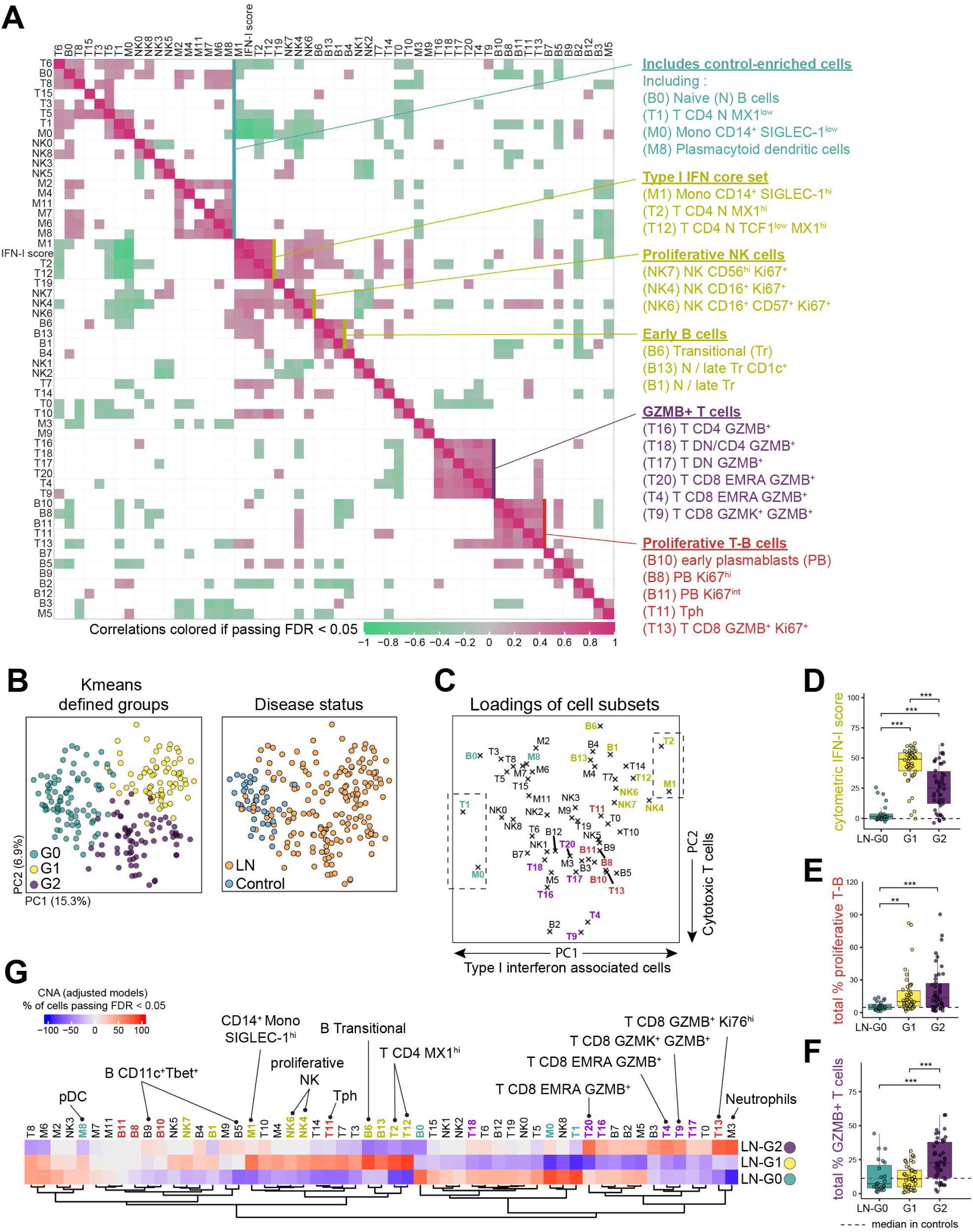
Unbiased analysis of blood immunophenotype identifies groups of lupus nephritis (LN) patients with distinct immune cell signatures. (**A**) Correlation plot organized by hierarchical clustering showing Spearman’s rho correlation coefficient between 55 immune cell clusters as well as type I interferon (IFN) cytometric signature in 115 patients with LN. (**B**) Main principal components distribution of 224 samples (39 controls, 115 baseline LN samples, 39 LN samples from week-12 and 31 LN samples from week-52) based on the proportion of 55 immune cell subsets and colored by K-means-defined groups (left panel) of by disease status (right panel). (**C**) Immune cell subset loading on the first two principal components. (**D**-**F**) Comparison of key immune cell signatures between each LN groups. Statistical significant was determined using Kruskal-Wallis test followed by Bonferroni correction for multiple testing. (**G**) Heatmap of CNA associations of specific immune cell subsets with each LN groups relative to the two other groups. Colors represent the percentage of cells passing FDR < 0.05, as either enriched (red) or depleted (blue). The colors associated with selected cell clusters and groups of cell clusters (green, yellow, purple and red) are consistent through (**A**-**G**).

We observed two other distinct sets of co-abundant immune cell clusters. One set included Tph (T11), activated memory B/early plasmablasts (B10), plasmablasts (B8, B11), and CD8 GZMB^+^ Ki67^+^ (T13) cells, comprising a set of T cells and B cells with a shared proliferative state (Ki67^+^), consistent with a prior study of patients with newly diagnosed SLE(61) (**Figure 3A**). A second, distinct set of co-correlated T cells was unified by their expression of granzyme B, including CD8 (T4, T9, T20), CD4, and DN (T16, T17, T18) T cells (**Figure 3A**). The T13 CD8 GZMB^+^ Ki67^+^ cluster strongly correlated with both the proliferative and the granzyme B^+^ cell sets and also correlated with IFN score. However, non-proliferating granzyme B^+^ T cells did not correlate with type I IFN signature (**Figure 3A**), suggesting that an IFN signature and granzyme B expression represent two different axes of immune activation. Together, these results enabled visualization of distinct sets of highly correlated immune cell populations that are unified by specific functional features, providing a framework with which to interpret clinical associations of immune cell clusters in patients with LN.

### Blood immunophenotype identifies three subgroups of patients with LN

As a second step to this unbiased approach, we evaluated whether the composition of circulating immune cell populations could distinguish groups of patients with LN. Here we applied a K-means clustering approach including all LN and control samples stained with 4 panels (n = 224), based on the scaled proportions of 55 immune cell subsets. We identified three groups of samples presenting distinct patterns, indicating potential patient subgroups. We confirmed > 80% stability of our clustering results by repeating 1000 K-means clustering on a resampled dataset without replacement (**Supplemental Figure 7A**). PCA and UMAP visualization confirmed a distinct distribution of these three groups (**Figure 3B, Supplemental Figure 7B**). Group G0 included all the controls (n=39) and a minority of patients with LN (n=23), indicating that these patients with LN had a control-like immunophenotype (**Figure 3B**). The two other groups, G1 and G2, each included 46 patients with LN at baseline.

To understand the main differences observed between these subgroups, we first examined the cell subset loadings on the first two PCs. We noted that both extremes of PC1 were marked by cells correlated with type I IFN score, suggesting that IFN response represented the main axis of variation in blood immunophenotypes (**Figure 3C**). Type I IFN-correlated subsets, including proliferating NK cells and transitional B cells, were significantly enriched in G1 compared to the other groups (**Supplemental Figures 7C-D**). Comparing the type I IFN score between the 3 groups of LN, G0 displayed the lowest values compared to the other groups and was not significantly different from controls. Group G1 was marked overall by the highest type I IFN values, whereas group G2 showed and intermediate level on average and the highest variability across patients (**Figure 3D**).

B and T cells characterized by a proliferative state (B8, B10, B11, T11 and T13) were enriched in both groups G1 and G2 compared to G0. In contrast, group G2 was marked by enrichment in granzyme B^+^ cell subsets (**Figures 3C and 3E, Supplemental Figure 7E**). The negative axis of PC2 appeared predominantly driven by T cells with expression of granzyme B (T4, T9, T16, T17, T18, T20 and T13), and the proportion of granzyme B^+^ T cells was increased in G2 compared to G0 and G1 (**Figures 3C and 3F, Supplemental Figure 7F**). As a complementary approach to the PC loadings, we used CNA to evaluate cells associated with G0, G1, and G2, adjusting for demographic characteristics, history of previous renal biopsy, prednisone dose and mycophenolate mofetil (MMF) use. This analysis confirmed the strong enrichment in granzyme B^+^ T cells in group G2. In addition, G2 was also characterized by an expansion of the main neutrophil cluster (M3) in the multivariable CNA model. B cells subsets enriched in G2 included “altered” naive CD19^lo^ CD21^lo^ (B3), and to a lesser extent CD11c^+^ (B5, B9) clusters (**Figure 3G**). Thus, this clustering approach distinguished 2 subsets of immunologically active patients with LN: one group (G1) with a prominent IFN signature, and a second group (G2) with a prominent expansion of granzyme B expressing lymphocytes.

### Baseline blood-defined LN groups have distinct renal pathology and outcome

We next explored whether these groups of patients with LN differed in relevant renal features. Patients in the group G0 (‘control-like’) had the lowest renal activity index scores, while the group G2 (‘cytotoxic T-enriched’) had the highest renal activity scores (**Figure 4A, Supplemental Table 3**). Consistent with the low activity index, the group G0 was more frequently classified as pure membranous, whereas G1 and G2 were more frequently classified as proliferative (85% in G2, 65% in G1, 35% in G0, p < 0.001). Detailed analysis of NIH renal activity index components revealed a significant correlation between G2 and endocapillary hypercellularity, active crescents, and fibrinoid necrosis, further underscoring active and more severe glomerular lesions in group G2 (**Figure 4B**). In addition, G2 correlated positively with the interstitial activity subscore, although results were at the limit of the FDR threshold of < 0.05 (p=0.01, FDR=0.05). Consistent with a more severe disease in the group G2 compared to G1 at baseline, those in the former group had increased serum creatinine (p=0.04), UPCR (p=0.009), and higher prednisone dose (p<0.001). In contrast, extrarenal manifestations and serologic measures (positive anti-dsDNA and/or low complement) were not significantly different between G1 and G2 (**Supplemental Table 3**). Adjusting for demographic variables, history of previous biopsy and baseline prednisone dose, the group G2 remained significantly associated with increased renal activity indices (beta coefficient [95%CI] = 2.6 [0.8-4.5], p.adj = 0.006) and proliferative class (OR [95%CI] = 4.2 [1.4-14.1], p.adj = 0.015) compared to the other groups (**Figure 4C**).

**Figure 4.**
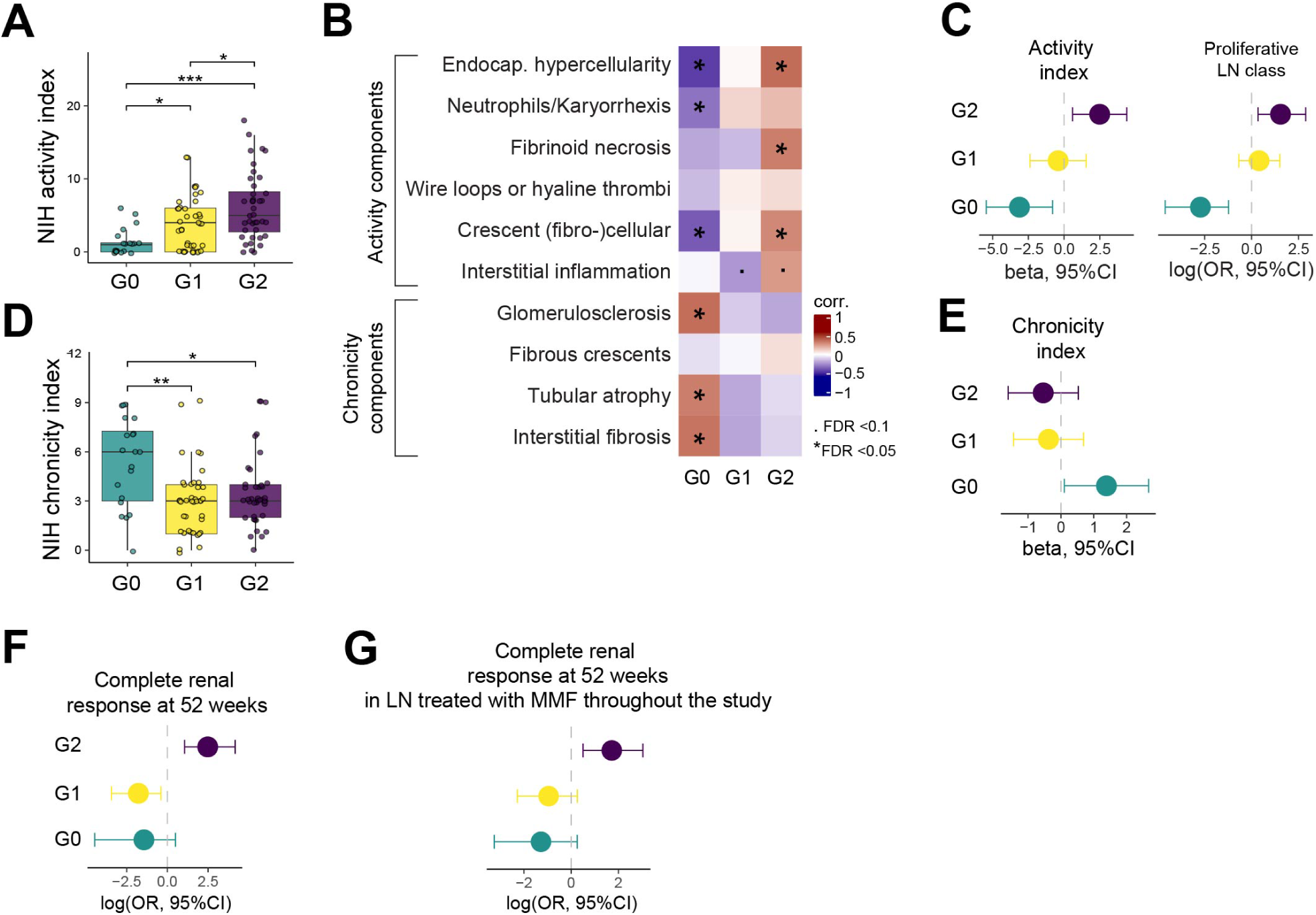
Blood-defined LN groups are associated with renal pathology and outcome. (**A**) Comparison of available NIH activity index between K-means defined groups (20 G0, 40 G1 and 40 G2) of patients with LN. Statistical significant was determined using Kruskal-Wallis with Dunn’s multiple comparisons. (**B**) Heatmap of Spearman’s rho correlation coefficient between each group against others and each subscores of the NIH activity and chronicity indices, adjusting for multiple testing (FDR < 0.05 and < 0.1 are indicated). (**C**) Multivariable models testing one-vs-rest group association with activity index (linear model) and proliferative class relative to membranous (logistic regression), accounting for demographic variables, history of previous biopsy and prednisone dose. (**D**) Comparison of available NIH chronicity index between K-means defined groups (n=20 G0, n=40 G1 and n=40 G2) of patients with LN. Statistical significance was determined using Kruskal-Wallis with Dunn’s multiple comparisons. (**E**) Multivariable model testing one-vs-rest group association with chronicity index (linear model), accounting for demographic variables, history of previous biopsy and prednisone dose. (**F**) Multivariable model testing one-vs-rest group association with response status (non-complete response vs complete response), accounting for demographic variables, history of previous biopsy and prednisone dose. (**G**) Univariate model testing one-vs-rest group association with response status in patients treated with MMF throughout the study.

Surprisingly, the group G0 had the highest NIH chronicity index and subscores, whereas G1 and G2 were not significantly different (**Figure 4D, Supplemental Table 3**). We observed that patients in G0 were older and more frequently had a history a previous biopsy (**Supplemental Table 3**). Examining the detailed classes in previous biopsies, we noted a significant increase in past proliferative (III, IV +/- V) disease in the group G0 compared to G2 (65% G0, 59% G1 and 33% G2, overall p=0.01, G0 v G2 p = 0.02). The majority (73%) of the patients with a membranous class in the group G0 had prior history of proliferative class, a histologic conversion previously reported(62). Adjusting for demographic variables, history of previous proliferative disease and baseline prednisone dose, G0 remained significantly associated with chronicity compared to the other groups, although to a limited extent (beta coefficient [95% CI] = 1.3 [0.04-2.5], p.adj = 0.044, **Figure 4E**). Together these results suggest that G0 identified a group of patients with little evidence of immune activation in blood, which is associated with minimal current renal activity, yet who may have previously had severe LN that evolved towards chronic renal damage.

We then asked whether these groups of patients had different renal outcomes. In the 88 patients with a defined renal response at week 52 (baseline UPCR ≥ 1), we observed a significant difference across groups (p=0.04) and increased renal response in G2, compared to G1 (p=0.03). Adjusting for demographic variables, history of previous biopsy, and baseline prednisone dose, the group G2 had an increased likelihood of complete response at one year (OR [95%CI] = 8.5 [2.2-40.5], p.adj = 0.003, **Figure 4F**). Given the variability in treatment received, we additionally tested the likelihood of response in patients treated with MMF throughout the study (MMF use at week 26 and 52: 52% in G0, 61% in G1 and 50% in G2), and observed a persistent association between G2 and complete response compared to others (OR [95%CI] = 5.6 [1.7-20.8], p = 0.007, **Figure 4G**). These results indicate that G2, characterized by increased proliferative and non-proliferative granzyme B^+^ T cells, neutrophils and B cell alterations, identifies a group of patients with severe, active disease but with increased likelihood of response to standard of care, including to an MMF-based regimen.

### Baseline blood-defined LN groups differ in kidney immune cell infiltrates and urine profiles

Given the differences in disease activity and outcome between these groups and previous reports associating type I IFN signaling with poorer renal response(6, 7), we hypothesized that the groups of patients with LN defined by blood immunophenotyping would show differences in the composition of the immune infiltrate within the kidney and type I IFN immune response gene expression. We leveraged scRNA-seq data generated on cells from kidney biopsies of 101 patients from the AMP phase II study(37, 63, 64). Given the prominent cytotoxic CD8 T cell features of G2 in the blood, as well as the previously recognized presence of CD8 T cell infiltrates in the periglomerular and interstitial space in proliferative LN kidneys(20, 65, 66), we then asked if CD8 T cell accumulation in the kidney differed in the blood-defined LN patient groups. Indeed, G2 patients showed an enrichment in granzyme B^+^ CD8 T cells, as well as a subset of granzyme K^+^ cells, in the kidney compared to the other LN groups (**Figures 5A-C**). In contrast, G2 patients were not associated with populations of CD8^+^ T cells enriched in *ITGA1, ZNF683* and *XCL1,* and depleted in *KLF2* expression, features suggestive of resident memory T cells(67, 68), which are also present in control kidneys(5) (**Figure 5A**). In addition, we observed differences in subsets of CD4 T cells in the kidney, which were mostly enriched in G0 and reduced in G2 (**Figure 5A-B**). Using a 21-gene type I IFN list(13, 69), we found that G1 patients had the highest IFN type I score in kidney immune cells, while G0 the lowest values, mirroring the pattern seen in the blood (**Figure 5D**). In total, these results indicate that immunoprofiling based on blood immune cells can identify subsets of patients with LN that differ in their renal immune cell infiltrates, with specific, direct relationships between immunophenotypic features in blood and those in kidney.

**Figure 5.**
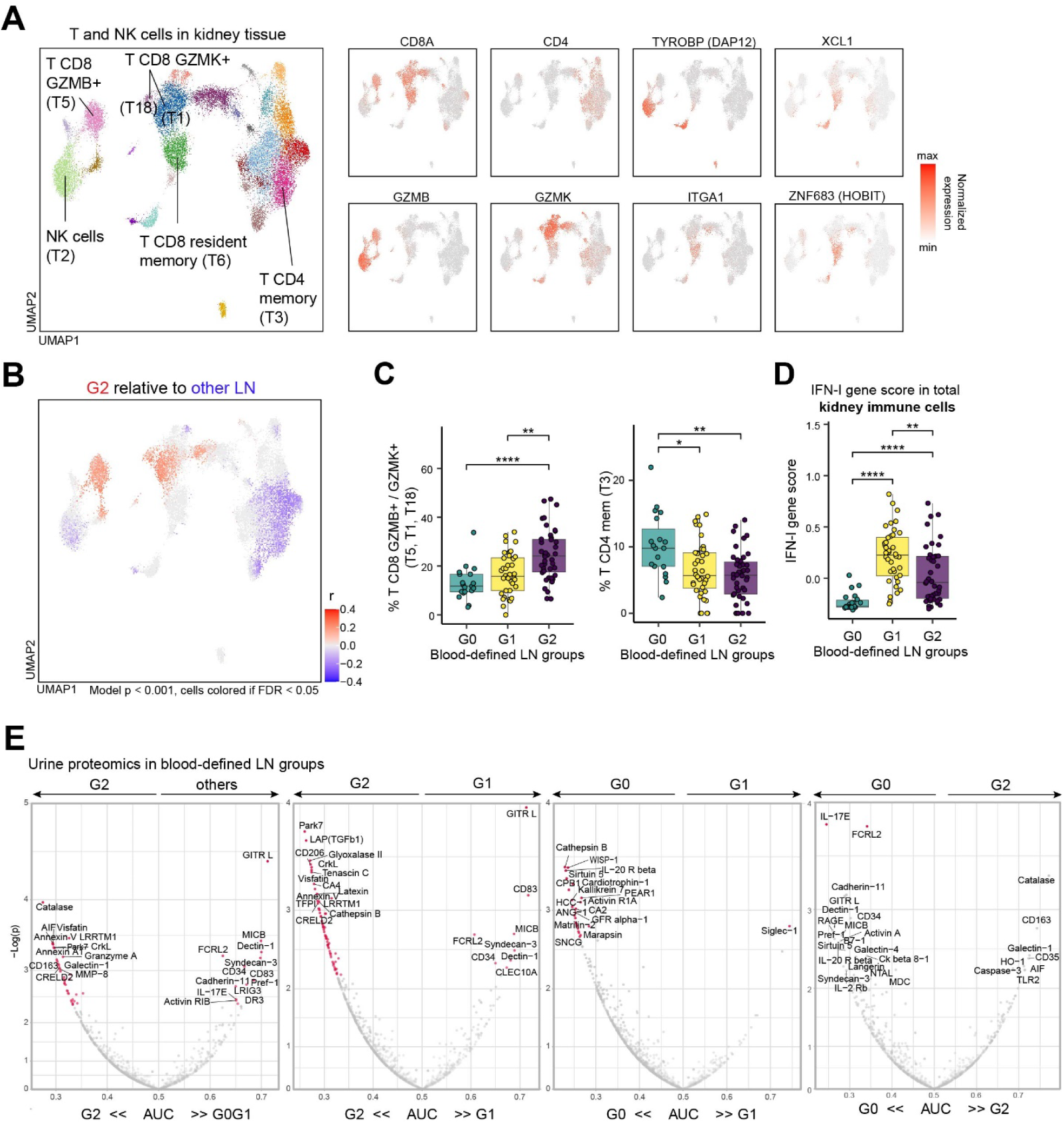
Blood-defined LN groups are associated with specific immune cell infiltration in the kidney and translate to urinary markers of inflammation. (**A**) Distribution of T cells and NK cells and selected gene expression by scRNA seq analysis of kidney tissue from patients with LN. (**B**) CNA association of T-NK cells with LN group G2 relative to others (G0 and G1) applied to paired kidney tissue samples. (**C**) Comparison of the proportions of CNA-identified granzyme B+ and a subset of granzyme K+ immune cell subsets, and T CD4 memory cells in the kidney tissue between groups. (**D**) Comparison of type I interferon (IFN) gene signature (21-gene list previously reported) in all immune cells from the kidney tissue between blood-defined LN groups. Statistical significant was determined using Kruskal-Wallis with Dunn’s multiple comparisons in (C and D), *p < 0.05, ** p < 0.01, ***p < 0.001. (**E**) Comparison of each urine proteine abundance between specified groups, as displayed by the performance of classification using the AUC (area under the curve). Dots colored in pink passed FDR threshold of 0.10, and labels were assigned to the top associated proteins. Statistical significance was determined using Wilcoxon rank sum test.

To further extend the association between blood immune profiles and kidney injury, we compared the urine proteomic profiles obtained from paired samples (n=115), previously generated using the Raybiotech kiloplex technology and published separately(35). After adjusting for FDR < 0.10, we observed significant differences between groups (**Figure 5E**). The group G2 was enriched in granzyme A and with proteins previously associated with increased NIH activity and proliferative LN class, such as Catalase and CD163(35, 70) (**Figure 5E**). Compared to the group G1, G2 had increased secretion of several proteins, including TGFβ1, which had been reported to be enriched in kidney tissue during proliferative LN flares and was predictive of complete renal response(7) (**Figure 5E**). In contrast, G1 was enriched in GITR-L and CD83, proteins expressed by antigen presenting cells, when compared to G2 and was significantly enriched in Siglec-1 when compared to G0 (**Figure 5E**). Together these results highlighted the association between blood immune alterations and kidney inflammatory process in patients with LN.

### Proliferating T-B cells decrease over time in MMF-based treatment responders

Given our results indicating that specific blood immune cell profiles associated with renal activity and outcome, we next aimed to understand how treatment may impact these signatures at baseline, in the context of a clinically diverse LN cohort, and longitudinally. We first used a linear model with penalization at baseline to simultaneously examine the importance of demographic features, renal and extrarenal characteristics, as well as medications, in accounting for the variation in immune cell signatures. We focused this analysis on 5 key blood immune signatures identified in the supervised and unsupervised analyses: type I IFN, proliferative T-B cells, non-proliferative granzyme B^+^ T cells, altered naïve B cells, and low-density neutrophils (**Figures 1-3**). Consistent with preceding analyses, the NIH renal activity index independently positively correlated with an increase in T-B proliferative cells (including T CD8 GZMB^+^ Ki67^+^ cells), low-density neutrophils and altered naïve B cells, whereas non-proliferative cytotoxic T cells were positively associated only with the interstitial component of the activity index. Type I IFN was not associated with renal activity (**Figure 6A**). Prednisone dose at baseline (excluding high intravenous dose of 500-1000mg) was an additional independent factor associated with increased low-density neutrophils, whereas MMF was independently associated with decreased proliferative T-B cells (**Figure 6A-B**).

**Figure 6.**
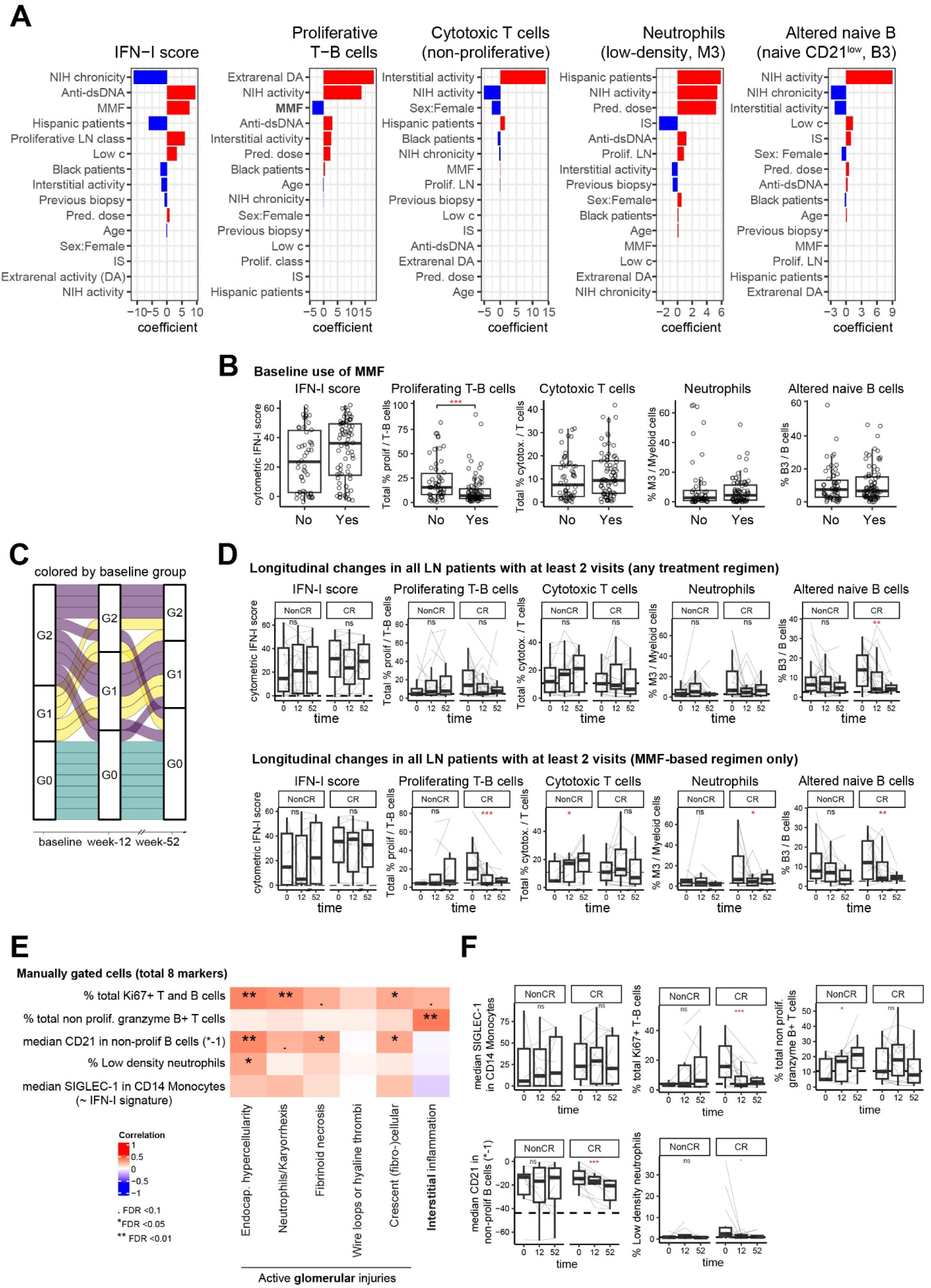
Immune cellular signatures heterogeneity across and within patients with LN. (**A**) Importance and direction of the effect of demographic, clinical and renal characteristics on immune cell signatures. The coefficient were defined using a linear model with an elastic net penalization using a 10-fold cross-validation, with the cellular signature as a response variable and the y-axis variables as the predictor variables. (**B**) Comparison of immune cell signatures between treated (Yes) or not treated (No) with MMF at baseline. Statistical significance was determined using Wilcoxon rank sum test test, ***p<0.001. (**C**) Changes in blood-defined group membership over time. Each band represent a patient, and each patient is colored by the baseline group membership. All patients with LN with samples at three timepoints and samples stained with all four panels were included in this analysis. (**D**) Longitudinal changes in immune cell signatures, stratified by response status (NonCR include none and partial responders, CR includes all complete responders). The upper panels include all patients with LN with at least 2 timepoints. The lower panels include patients with LN who were treated with MMF throughout the study. Statistical significance was determined using a mixed effect model including patient as a random effect. *p<0.05, **p<0.01, ***p<0.001. (**E**) Correlations between simplified immune cell signatures and NIH activity subscores at baseline. Statistical significance was determined using Spearman’s rho correlation followed by FDR correction for multiple testing. For visualization purposes we showed the inversed value of median CD21 in non-proliferative B cells (written as *-1). (**F**) Longitudinal changes in simplified immune cell signatures in patients treated with an MMF based therapy. For visualization purposes we showed the inversed value of median CD21 in non-proliferative B cells (written as *-1). Statistical significance was determined using a mixed effect model including patient as a random effect. p = 0.05, *p<0.05, **p<0.01, ***p<0.001.

We then examined the group changes longitudinally in these signatures. Immunologically inactive patients (group G0) at baseline uniformly remained stable over time (**Figure 6C**). In contrast, the majority of patients who started in G2 changed their group over time, including some transitioning to G0, suggesting that the cellular features of G2 are dynamic and can change with clinical course and treatment. Detailed analysis of immune signatures showed that the proportion of altered naïve B cells decreased over time in complete responders but not in partial and non-responders (**Figure 6D**), while immune features of IFN score, proliferating T-B cells, and cytotoxic T cells did not change consistently in either responders or non-responders (**Figure 6D**). However, the decreased trend observed in proliferative T-B cells was significant in responders when including only patients with an MMF-based regimen, suggesting a drug effect (**Figure 6D**). In contrast, the type I IFN signature did not significantly change over time, including in patients treated with MMF, supporting previous reports in general SLE(23, 48) that this pathway is not robustly altered by standard of care LN regimens.

Finally, we confirmed that these 5 main cellular features could be captured using a simplified manual gating strategy using a limited number of canonical markers (**Supplemental Figure 8**). To minimize the number of markers, we measured the expression of CD21 in non-proliferative B cells and for visualization purposes (**Figures 6E-F**) we showed the inversed value, for consistency with previous graphs. Overall, this manual gating strategy, applied to the total dataset of live cells (25-30^10^6^), reproduced the key patterns of cellular features associated with renal disease activity and longitudinal changes, including the decrease in proliferative T-B cells and the increase in CD21 expression in non-proliferative B cells over time, supporting that these populations can readily be identified (**Figure 6E,F**).

## Discussion

Here, we have used blood cell immunoprofiling to identify immunologically distinct groups of patients with LN that differ in renal inflammation and outcome in a large cohort of patients who underwent a clinically-indicated renal biopsy. Unsupervised modeling approaches visualized co-abundant cell populations and stratified patients with LN into 3 groups: an immunologically inactive group (G0), a predominant type I IFN group (G1), and a predominant cytotoxic T cell group (G2). Both immunologically active groups (G1, G2) show increased proportions of proliferating T cells and B cells, yet these 2 groups differ with much higher IFN scores in G1 and increased proportions of cytotoxic T cells, low density neutrophils, and CD21^low^ altered naïve B cells in G2. These results reveal specific features that distinguish the immunologic heterogeneity among patients with LN, highlighting in particular that features of an activated cellular response characterizing the group G2 can be separated from a high IFN response. Further, a separate group of patients with biopsy-demonstrated LN, patients in G0, show little evidence of immune activation, with immunologic profiles that are indistinguishable from control patients.

LN patient groups distinguished by this blood immunophenotyping approach differed in renal injury and renal immune cell infiltration, indicating that blood profiling can reflect relevant intra-renal features. Group G0 patients, who appeared immunologically quiet by immunophenotyping, also showed little renal activity, although had the highest scores for chronic renal damage. The history of prior episodes of proliferative nephritis in these patients suggests that these patients may have proteinuria due to chronic renal damage but little ongoing active inflammation. Group G1 patients showed the highest type I IFN scores in the blood and associated with an increase in circulating proliferative NK cells and increased transitional B cells, and correspondingly showed the highest expression of IFN response in immune cells from kidney. IFN responses may alter the functions of many kidney-infiltrating immune cells as well as kidney parenchymal cells(71–74). The high IFN signature in blood cells, kidney cells, and in urine of these patients suggests a globally high IFN response, which can be robustly captured by blood cytometry, consistent with prior reports(43, 75).

The G2 group, characterized by the expansion of granzyme B^+^ T cells, showed the most active proliferative renal disease. This is consistent with prior reports of expanded effector memory T CD8 cells in the blood of SLE patients with more active disease(76–78). Our detailed profiling approach revealed a set of co-correlated granzyme B^+^ T cell populations, including subsets of CD8 T effector memory and terminally differentiated effector memory populations, proliferating CD8 T cells, CD4 T cells and CD8^-^CD4^-^ T cells, associated with this group of patients with LN. We further demonstrated that this G2 patient group has a concomitant enrichment in paired kidney samples of GZMB^+^ and GZMK^+^ T cell subsets, a major driver of tissue inflammation in autoimmune diseases(79). A pathogenic role for CD8 T cells in patients with LN has been previously suggested by the enrichment in cytotoxic and proliferating CD8 T cells and production of inflammatory chemokines and interferon-γ(5, 9), a cytokine implicated in the pathogenesis on LN disease(80) in kidneys of patients with LN(65, 66). A direct connection between CD8 T cells in blood and kidney is suggested by the description of shared CD8 T cell clones between these sites in patients with LN(66). Interestingly, we identified different correlations involving specific T cell phenotypes: circulating proliferating cells (including granzyme B^+^ T cells) correlated with glomerular pathology, whereas non-proliferating granzyme B^+^ T cells correlated with interstitial renal activity.

Notably, the G2 group with a predominant cytotoxic T signature at baseline showed an increased likelihood of response to standard of care at one year compared to the other groups. This is somewhat surprising because expansion of effector CD8 T cells in SLE patients and infiltrating CD8 T cells in LN have been associated with more severe and refractory disease(65, 66, 78). In addition, tubulointerstitial activity has been previously associated with poorer outcome(81). However, by assessing the immune alterations comprehensively, we also found that this group of patients had a distinct urine proteomic profile, suggesting broader immune differences between groups. We hypothesize that these features reflect aspects of immune cell activation that can effectively be suppressed by current immunosuppressive therapies, such as MMF, which was associated with reduction in low density neutrophils and CD21^lo^ naïve B cells populations in this cohort. Additionally, the two other groups of patients identified were either marked by high fibrotic histologic lesions (G0) or high type I interferon responses (G1), both signatures previously associated with poorer outcome in the AMP phase I study(6). In addition, we showed that the type I IFN pathway was not reduced by standard of care immunosuppressive therapy, potentially contributing to the low renal response rate in this group. The particularly high expression of type I IFN, contrasting with the variability in the group G2, suggests that considering the intensity of IFN response and combining it with other immune cell signatures might help stratify patients more effectively than the commonly used division of IFN-high versus IFN-low patients. Further studies will help define if patients with a disease characterized by high type I IFN benefit from therapies that effectively disrupt this pathway.

Our study has limitations. Although we examined the blood immunophenotype using 4 panels with 48 proteins each, other relevant signals might be present and could help further determine key cellular features associated with the immune cells enriched in the different groups of patients, including cytokine profiles enriched in the different groups. Ongoing work, as part of the AMP phase II study, on transcriptomic profile of immune and resident kidney cells, as well as blood immune cells will provide more detailed blood-tissue comparison including comprehensive signals potentially further distinguishing these groups of patients with LN. Secondly, controls were not well-matched to our cohort of patients with LN. To account for demographic differences, we systematically analyzed our data in univariate and multivariate models adjusting for age, sex, ethnicity and race. Thirdly, this study was limited by the lack of standardized therapy. Despite these challenges, we were able to identify immune cell changes that were either associated with or independent of the immunosuppressive therapies. Detailed assessment in a randomized trial with standardized therapy will further help understand the immune changes associated with the heterogeneity of response to specific therapies.

In summary, our study highlights the power of blood immunophenotyping to define subsets of patients with LN with distinct patterns of immune activation that reflect ongoing renal inflammation and injury, which provides critical evidence for future personalized treatment. This work indicates a valuable role for blood cellular biomarkers in evaluating specific dimensions of immune activity of patients with LN to gain insight into the likelihood of response to standard immunosuppressive LN therapies.

## Method

### Study population

SLE patients 16 years or older were recruited between January 2016 and May 2021 at 14 clinical sites across the United States as part of the Accelerating Medicines Partnership (AMP) RA/SLE Phase II study(37, 82) if 1) they fulfilled American College of Rheumatology(83) or Systemic Lupus International Collaborating Clinics classification criteria(84) for SLE, 2) had a clinically-indicated renal biopsy for a UPCR > 0.5(16, 17) and 3) had biopsy-confirmed lupus nephritis of class III, IV and/or V, based on the International Society of Nephrology/Renal Pathology Society (ISN/RPS)(15). Patients with a history of renal transplantation, with a medical condition considered at risk of participating in this study by the investigator, who was pregnant at the time of study entry, or who had recently been exposed to rituximab (within the past six months) were excluded. Healthy controls were recruited from 2 of the clinical sites.

### Data and sample collection

Patient characteristics and clinical assessment were collected at each site, including demographic characteristics (age, sex, ethnicity and race), history of previous renal biopsy, SLE disease activity measured by the hybrid Safety of Estrogens in Lupus Erythematosus National Assessment (SELENA)-SLE Disease Activity Index (SLEDAI) (urine protein to creatinine ratio > 0.5 always counted)(85, 86), anti-dsDNA antibody status (according to the upper limit of the norm of each site laboratory), C3 and C4 complement values, serum creatinine, estimated glomerular filtration rate, UPCR, and medications received during the study. Patients were treated during the study at the discretion of their physicians based on standard of care. Renal response to treatment was determined clinically at week 52 as complete, partial, or none, adapted from the ACCESS trial definitions(87), and as reported(35), in patients with a baseline UPCR ratio ≥ 1. Briefly, complete response (CR) was defined by UPCR <0.5, normal serum creatinine (≤ 1.3 mg/dL) or, if abnormal, <125% of baseline, and prednisone ≤10mg/day. Partial response (PR) was defined by >50% reduction in UPCR without meeting UPCR criterion for CR, normal creatinine (≤ 1.3 mg/dL) or, if abnormal, ≤ 125% of baseline, and prednisone dose ≤ 15 mg/day.

Renal biopsy sections were classified and scored according to the ISN/RPS classification as proliferative (class III, IV +/- V) or membranous (class V), and the NIH activity and chronicity indices(15). Initial assessment was performed by a renal pathologist at each site, followed by a central read by two independent renal pathologists for 101 biopsies. Central reads included a detailed subscoring of each component of the NIH activity and chronicity indices. Scores from central reads were used for further analysis if available, and total scores from the individual site reads were used otherwise.

Blood samples were collected from patients with LN and controls at baseline (week 0), and a subset of patients with LN were recollected at weeks 12 and 52. PBMCs were isolated from blood samples and cryopreserved at each site as previously described(32). Cryopreserved PBMC were shipped to the central AMP RA/SLE Biorepository, Oklahoma Medical Research Foundation Biorepository, for storage until sample collection was complete. A total of 275 samples were collected from 152 patients with LN and 40 controls (**Supplemental Figure 1A**).

### Sample staining and data acquisition by mass cytometry

Cryopreserved PBMC samples were sent to the Brigham and Women’s Hospital CyTOF Antibody Resource and Core. Samples were then randomly distributed across 23 staining and acquisition days (23 batches) at 20 samples per day (**Supplemental Figure 1A**). Samples were thawed, stained and mass cytometry data was generated according to the AMP RA/SLE mass cytometry processing workflow, and as previously reported(88). As described, samples were thawed in a 37 °C water bath for 3 minutes and then mixed with 37 °C thawing media containing: RPMI Medium 1640 (Life Technologies #11875-085) supplemented with 5% heat-inactivated fetal bovine serum (Life Technologies #16000044), 1 mM GlutaMAX (Life Technologies #35050079), antibiotic-antimycotic (Life Technologies #15240062), 2 mM MEM non-essential amino acids (Life Technologies #11140050), 10 mM HEPES (Life Technologies #15630080), 2.5 x 10^-5^ M 2-mercaptoethanol (Sigma-Aldrich #M3148), 20 units/mL sodium heparin (Sigma-Aldrich #H3393), and 25 units/mL benzonase nuclease (Sigma-Aldrich #E1014). 100 μL aliquots of each sample post-thaw were mixed with PBS (Life Technologies #10010023) at a 1:1 ratio to be counted by flow cytometry.

Each sample was then split into 4 to be stained with 4 different panels including lineage and specific markers dedicated to examining B, T, myeloid, and NK cells (**Supplemental Figure 1A**). Overall, between 0.5 – 1.0 x 10^6^ cells were stained per panel for each sample. If the sample before splitting included <2×10^6^, a panel priority was used: B panel > T panel > Myeloid panel > NK panel. The list of markers is detailed in **Supplemental Table 4**. All samples were transferred to a polypropylene plate (Corning #3365) to be stained at room temperature for the rest of the experiment. The samples were spun down and aspirated. Rhodium viability staining reagent (Standard BioTools #201103B) was diluted at 1:1000 and added for five minutes. 16% stock paraformaldehyde (Fisher Scientific #O4042-500) was diluted to 0.4% in PBS and added to the samples for five minutes. After centrifugation and aspiration, Human TruStain FcX Fc receptor blocking reagent (BioLegend #422302) was used at a 1:100 dilution in cell staining buffer (CSB) (PBS with 2.5 g bovine serum albumin [Sigma Aldrich #A3059] and 100 mg of sodium azide [Sigma Aldrich #71289]) for 10 minutes followed by incubation with conjugated surface antibodies (each marker was used at a 1:100 dilution in CSB, unless stated otherwise) for 30 minutes. The Harvard Medical Area CyTOF Antibody Resource and Core (Boston, MA) prepared and validated all antibodies. After centrifugation, samples were resuspended with culture media. 16% stock paraformaldehyde (Fisher Scientific #O4042-500) dissolved in PBS was used at a final concentration of 4% for 10 minutes to fix the samples before permeabilization with the FoxP3/Transcription Factor Staining Buffer Set (ThermoFisher Scientific #00-5523-00). The samples were incubated with SCN-EDTA coupled palladium barcoding reagents for 15 minutes, followed by incubation with Heparin (Sigma-Aldrich #H3149-100KU) diluted 1:10 in PBS. Samples were combined and filtered in a polypropylene tube fitted with a 40µm filter cap. Conjugated intracellular antibodies were added into each tube and incubated for 30 minutes. Cells were then fixed with 4% paraformaldehyde for 10 minutes. DNA was labeled for 20 minutes to identify single-cell events with an 18.75 μM iridium intercalator solution (Standard BioTools #201192B). Samples were subsequently washed and reconstituted in Cell Acquisition Solution (CAS) (Standard BioTools #201240) in the presence of EQ Four Element Calibration beads (Standard BioTools #201078) at a final concentration of 1×10^6^ cells/mL. Samples were acquired on a Helios CyTOF Mass Cytometer (Standard BioTools).

### Mass cytometry data processing and quality control

The raw FCS files were normalized to reduce signal deviation between samples during multi-day batch acquisitions, based on the bead standard normalization method established by Finck et al. (89). The normalized files were then compensated with a panel-specific spillover matrix to subtract cross-contaminating signals(90). These compensated files were then deconvoluted into individual sample files using a single cell-based debarcoding algorithm established by Zunder et al(91). Normalized and debarcoded FCS files were then uploaded to OMIQ software from Dotmatics (www.omiq.ai, www.dotmatics.com), as previously reported(88).

Manual gating on Gaussian parameters and DNA was applied to remove debris, outlier events, and doublets. Singlet live cells were then identified by gating on bead-negative and Rhodium-negative cell events. The data was further evaluated for quality control by examining the proportion of viable cells, the presence and frequency of major cell types (B, T, myeloid, and NK) and the distribution of samples using unsupervised clustering with opt-SNE dimensionality reduction and PARC clustering(92), available in the OMIQ software from Dotmatics (www.omiq.ai, www.dotmatics.com). A total of 8 samples were excluded from further analysis due to one or several quality issues: cell viability < 50%, 0 B cells, and samples identified mostly (>90%) in one PARC cluster. Thus, further analysis included 264 samples from 145 patients with LN and 40 controls (**Supplemental Figure 1A**, **Supplemental Table 2**). In total, over 110 million cells were analyzable across panels, and the median cell counts per sample within each panel after quality control was above 100,000 cells (**Supplemental Figure 1B**).

### Data transformation and batch correction

The data generated from each panel was analyzed separately using the same strategy. After quality control filtering, FCS files including live singlet cells, were uploaded and read in R (R version 4.3.1) using the flowCore package (version 2.14.0) and we then applied arcsinh-transformation with a cofactor of 5 to the protein expression matrix, as described(93). We leveraged the single-cell transcriptomic analysis pipeline from the Seurat package (version 4.3.0)(94) for our workflow by transposing our transformed expression matrix and implementing it into a Seurat object with the corresponding metadata. To ensure equal representation of samples (**Supplemental Figure 1B**) and to save computational time and resources, we randomly downsampled each sample to 10,000 cells (if a sample had < 10,000 cells, all cells were kept), as downsampling does not affect the sensitivity of the clustering analysis(88).

All proteins in each panel were examined and included in further analysis, except for IgG and IgA in the B panel, due to an identified strong co-expression of both markers that was determined to be a technical limitation. We used principal component analysis (PCA) for dimensionality reduction (function RunPCA, set without approximation) from the Seurat package and then corrected batch effects with the function RunHarmony based on the Harmony(38) application programming interface implemented in the Seurat package, using the first 20 PCs (**Supplemental Figure 1C-D**). We quantified the integration of the batches using the function compute_lisi from the LISI package (version 1.0)(38) (**Supplemental Figure 1E**).

### Graph-based clustering, visualization of protein expression and cell subset annotation

After batch correction, we obateined a nearest-neighbor graph (function FindNeighbors) and clusters based on a Louvain-based algorithm (function FindCluster) from the Seurat package(94). Cells were then projected into two dimensions using the runUMAP function. We annotated each cluster as T, B, Myeloid or NK cells based on lineage markers in each panel. Clusters that were not attributed clearly to one of these cell types were labeled undetermined and were excluded from further analysis (< 0.1% of total live cells for each panel) (**Supplemental 1F-H, Supplemental Method**).

As a second step, we extracted the cell type of interest in the dedicated panel (e.g., T cells in the T panel) and re-clustered the cells by applying again the functions: RunPCA(), RunHarmony(), FindNeighbors(), FindCluster() and RunUMAP(). For the Louvain-based algorithm, we optimized resolution for each cell type (0.8 for B cells, 0.7 for T cells, 0.3 for Myeloid cells, 0.3 for NK cells), based on the cluster distribution and manual check of expression of critical proteins in each cluster to gain the biological interpretations that made the most sense. Cell-type specific clusters were labeled with a first letter corresponding to the panel they were extracted from, and a number based on the cluster size (e.g., the cluster “B0” corresponds to the largest cluster within B cells extracted from the B panel). Strategies for visualization are detailed in the **Supplemental Methods**.

### Association of cell subsets with disease status and clinical characteristics

We applied co-varying neighborhoods analysis (CNA)(41) using its r version (rcna package, version 0.0.99) implemented in the Seurat package (association.Seurat function). For each model, we examined and reported the global CNA p-value, which is defined through a permutation test, and further evaluated the cell-neighborhoods that passed a threshold of FDR<0.05. Local associations passing the FDR threshold were mapped in the UMAP space. Separately, we applied the mixed-effects association of single cells (MASC)(42) and examined the consistency between the results of the 2 approaches, by testing the correlation between the median cell correlation coefficient, defined by CNA, for each cluster, and the odds ratio defined by MASC (**Supplemental Methods**).

### Protein association with the dominant signal in CNA

We determined the correlation between NAM-PC1 coordinates (obtained from the CNA analysis) and each protein expression in the panel, at a single-cell level (**Figure 1D,E**). We obtained the spearman’s coefficient of correlation between NAM-PC1 and individual protein expression using the rcorr function from the package Hmisc. Results were validated visually by mapping the expression of the highest marker identified in the UMAP space in comparison to the mapping of CNA results.

### Cytometric type I interferon score calculation

We identified any proteins in our panels that had been previously and repeatedly used as a gene or a protein associated with type I interferon signaling(13, 25, 43, 75). We calculated the median expression of MX1 and ISG15 amongst total live cells, as the expression was broadly observed for both markers. In contrast we used the median expression of SIGLEC-1 amongst myeloid cells, as no other cell type expressed it (**Supplemental Figure 3E**). For each sample stained with these three panels, the values for each marker were standardized to the mean and standard deviation of the controls and then summed to obtain a score.

### Identification of circulating immune cell signatures

We extracted the proportions of each cell subsets among cell type for each sample and retained 55 cell subsets (=cell subset) with < 50% of zeros across samples. Including all baseline samples from patients with LN stained with four panels (n=115), we obtained the Spearman’s rho correlations coefficients and p values between all cell subsets and type I interferon score using the rcorr function from the Hmisc package (version 5.1-1) and the FDR using the p.adjust function from the stats package (version 4.3.1). Results were visualized using the R package corrplot (version 0.92) by applying hierarchical clustering with Ward distance method (**Figure 3A, Supplemental Figures 6A, Supplemental Methods**).

To identify the differential expression markers between two T CD4 naïve clusters associated with type I interferon signature, we used the limma package(95), a method used previously to evaluate differential expression in mass cytometry dataset(96), to compute the p values for each marker included in the T panel. After identifying TCF1 as the most differentially expressed marker, we examined it’s expression in CD4 naïve T cells after stimulation with CD3/CD28 with or without IFN beta, using a publicly available RNAseq dataset (GSE195541)(57).

### Identification of blood-defined immunophenotype groups

We then identified three immunophenotype groups based on K-means clustering on the 55 scaled cell subset proportions for each sample stained with four panels (including controls and LN at all timepoints) using the kmeans function from the stats package. To test for the stability of the clusters we repeating 1000 K-means clustering on a resampled dataset without replacement, using the ConsensusClusterPlus package (version 1.66.0)(97) and further examined the mean cluster consensus across samples for each k and obtained over 80% of stability with a k=3 (**Figure 3A**). We additionnally tested that the differences across groups were also reflected in the PCA and UMAP spaces using the umap function from the uwot package (version 0.1.16) (**Figure 3B, Supplemental Figure 7B**).

To identify the cells driving the differences between these groups, 1) we examined the cell subset loadings in the first 2 PCs (**Figure 3C**), 2) we run cell-type specific CNA comparing each group with the others including patients with LN at baseline and used a heatmap to combine the results (**Figure 3G**) and 3) we compared the proportions of a selection of cell subsets, either combined or individually (**Figure 3E-F, Supplemental Figure 7D-F**). For visualization of CNA results, we plotted the frequency of cells within each cell subset passing the FDR threshold of <0.05. All CNA models with cells passing FDR had a global p value of < 0.05.

### Association of blood-defined LN groups with T cell infiltrating cells and type I IFN score

As part of the AMP phase II study, human kidney biopsies were cryopreserved, thawed, dissociated, sequenced and processed as described(63). Following alignment, dimensionality reduction was achieved by identifying variable genes and running PCA, followed by batch correction using Harmony(38), graph-based clustering using Seurat (version 4.1.0)(94). T and NK cell clusters were identified based on the expression of known lineage markers (Al Souz *et al*., manuscript in preperation). Blood-defined groups were assigned to paired samples and CNA(41) was applied to examine the differences in cell neighborhoods between groups, in addition to examining the proportions of T cell clusters. To evaluate type I IFN signaling between groups, we used the AddModuleScore function (Seurat package) to define type I IFN score based on a previously described 21-gene list(13), across all immune cells in the kidney.

### Urine proteomic from paired patients with LN

The screening of urine proteomic was previously described and reported for patients with LN included in the AMP phase II study(35). Briefly, the screening was performed using an extended version of the Kiloplex Quantibody (RayBiotech) and the concentration of each analyte was normalized by urine creatinine to account for urine dilution. We compared the abundance of urine proteins in pg_protein_/mg_creatinine_ between blood-defined groups of patients with LN using a Wilcoxon rank sum test and adjusting for FDR, with a threshold of < 0.10 and < 0.25 as indicated.

### Statistics

In addition to the statistical testing mentioned above, we used non-parametric tests for cross-sectional univariate analysis at baseline: Wilcoxon rank sum test to compare continuous or ordinal variables between two groups, Kruskal-Wallis test followed by post hoc Dunn’s test to compare continuous or ordinal variables between more than two groups, or Spearman’s rho to assess the correlation between two continuous or ordinal variables (using wilcox.test, kruskal.test, dunn_test, cor.test with the method spearman functions in r). To compare categorical variables between groups, we used Fisher exact test and Chi-squared test depending on the size of the tested categorizes (using fisher.test and chisq.test functions in r). For multivariable models, we used linear regression models or generalized linear models for logistic regression (lm and glm function in r) to test for association with a continuous or categorical variable, respectively, including the covariate detailed in the results. To compare LN groups using logistic regression models, we applied a one-vs-rest strategy to examine the specific characteristic of one group compared to the others. To test for clinical and renal factors associated with the variation of selected cell subsets of interest, we applied a linear model with penalization using an elastic net regression (glmnet function used with the caret package) after 10 random repeats of a 10-fold cross-validation, including the cell proportion as the response variable and the clinical and renal variables as ‘predictor’ variables. All predictor variables were normalized to a fixed range between 0 and 1. For all above cross-sectional analysis, missing clinical data or samples were excluded from statistical testing. For longitudinal data, we used mixed effect models (lmer function from the lme4 package, version 1.1-34) to test for change over time by including each patient as a random factor to account for paired samples. For longitudinal data, we included all patients with a defined response status at 1 year who had at least two separate samples. All statistical tests were two-sided. For data visualization, we used the packages ggplot, ComplexHeatmap and corrplot. All analysis were performed on R version 4.3.1.

### Study approval

All participants provided written informed consent before study enrollment, and human study protocols were approved in accordance with the Declaration of Helsinki by the institutional review boards (IRBs) at each participating sites (detailed in **Supplemental Methods**).

## Data availability

Mass cytometry and clinical datasets will be made available on the ARK Portal upon publication. All analyses can be reproduced using the publicly available versions of the R packages outlined in the method and R scripts are available from the corresponding author on request.

## Authors contribution

AF, KP, MD, DW, DLK, KCK, RF, MB, PI, RC, DH, ESW, MAM, JG, JLB, FP-SMI, MW, MK, CP, JAJ, MAP, JPB recruited patients, obtained samples, and curated clinical data DAR, JAL, FZ, AF, TME, JG, PJH, MD, DW, DLK, KCK, RF, MB, PI, RC, DH, ESW, WA, MAM, JG, JLB, FP-S, MI, MW, MK, CCB, JBH, DSD, CP, MBB, JHA, SR, NH, JAJ, AD, MAP, JPB, and BD contributed to the processing of samples and design of the AMP SLE study JBH and DSD provided central histopathological read of the kidney biopsies AG, JK, JP, EM, KH, BH and, JAL designed and generated mass cytometry data AG and AH processed mass cytometry data with the support of TG and TS AH analyzed the data with the support of JI, FZ, MGA and DAR TS, RB and YC advised and supported on aspects of the data analysis TME, NH generated the scRNAseq data from kidney tissue and SR and AA processed the scRNAseq data AF analyzed the urine proteomic data AF, PJH, PI, MBB, JHA, SR, NH, JAJ, AD, MAP, JPB, BD, FZ, JAL, DAR provided disease clinical and immunology input AH and DAR wrote the initial draft DAR, JAL, and FZ supervised the research All authors participated in editing the final manuscript

## Acknowledgments

We would like to thank the patients who participated in this study, and the scientists and clinical sites in the Accelerating Medicines Partnerships in RA/SLE. This work was supported by the Accelerating Medicines Partnership® Rheumatoid Arthritis and Systemic Lupus Erythematosus (AMP® RA/SLE) Network^#^. AMP is a public-private partnership (AbbVie Inc., Arthritis Foundation, Bristol-Myers Squibb Company, Foundation for the National Institutes of Health, GlaxoSmithKline, Janssen Research and Development, LLC, Lupus Foundation of America, Lupus Research Alliance, Merck & Co., Inc., National Institute of Allergy and Infectious Diseases, National Institute of Arthritis and Musculoskeletal and Skin Diseases, Pfizer Inc., Rheumatology Research Foundation, Sanofi and Takeda Pharmaceuticals International, Inc.) created to develop new ways of identifying and validating promising biological targets for diagnostics and drug development Funding was provided through grants from the National Institutes of Health (UH2-AR067676, UH2-AR067677, UH2-AR067679, UH2-AR067681, UH2-AR067685, UH2-AR067688, UH2-AR067689, UH2-AR067690, UH2-AR067691, UH2-AR067694, and UM2-AR067678). This work was also conducted with support from UL1TR002541 award through Harvard Catalyst | The Harvard Clinical and Translational Science Center (National Center for Advancing Translational Sciences, National Institutes of Health) and financial contributions from Harvard University and its affiliated academic healthcare centers. Dr Horisberger fellowhip was supported by a grant from the Swiss National Science Foundation Early Postdoc.Mobility P2LAP3_199491 and by the Lausanne University Hospital in Switzerland. This work was also supported by the Arthritis National Research Foundation Award (to F.Z.).

## Notes

### Competing Interest Statement

D.A.R. and M.B.B. are co-inventors on patent on Tph cells as a biomarker of autoimmunity.

### Summary of Updates

Correction of figure labels in the text and reformatting for submission

