## Supplemental Tables for "Blood immunophenotyping identifies distinct kidney histopathology and outcomes in patients with lupus nephritis"

**Supplemental Table 1**. Baseline characteristics of all subjects included in the study and renal response to treatment at 52 weeks

| | **Lupus nephritis patients** | **Controls** | ***p value***[*] |
|---|---|---|---|
| **N** | **145** | **40** | |
| **Age, median (IQR)** | 33 (26-43) | 44 (27-60) | 0.02 |
| **Female** | 126 (87) | 28 (70) | 0.02 |
| **Ethnicity and race**[1] | | | |
| Hispanic or Latino | 45 (31) | 4/39 (10) | 0.008 |
| Black | 67/128 (52) | 10 (25) | 0.003 |
| **Previous renal biopsy** | 100 (69) | | |
| Previous LN class I or II | 12 (8) | | |
| Previous LN class III, IV and/or V | 88 (61) | | |
| **Current renal histologic ISN class** | | | |
| Proliferative (III or IV +/- V) | 102 (70) | | |
| Membranous (V) | 43 (30) | | |
| **Activity index[2], median (IQR)** | 4 (1-7) | | |
| **Chronicity index[2], median (IQR)** | 3 (2-5) | | |
| **Serum creatinine[3] mg/ml, median (IQR)** | 0.9 (0.7-1.2) | | |
| **UPCR, median (IQR)** | 2.0 (1.2-3.8) | | |
| **Presence of cSLEDAI extrarenal[4]** | 65/130 (50) | | |
| **Positive anti-dsDNA** | 96/142 (68) | | |
| **Low C3 and/or low C4** | 97/143 (68) | | |
| **Medication at baseline** | | | |
| Hydroxychloroquine | 121/144 (84) | | |
| Prednisone dose, med (IQR)[5] | 5 (0-25) | | |
| Any immunosuppressants[6] | 104/144 (75) | | |
| Mycophenolate | 82/144 (57) | | |
| Cyclophosphamide | 2/144 (1) | | |
| **Renal response at 52 weeks (n=112)**[7] | | | |
| CR, PR, NR | 31 (28), 27 (24), 54 (48) | | |

UPCR = urine protein/creatinine ratio. CR, PR, NR = complete, partial and non-responder. Data is presented as N (%) unless specified otherwise. The proportions, median and IQR are calculated on the total number of subjects in the group unless specified otherwise. *Chi-square, Fisher exact or Mann-Whitney tests were used when appropriate for comparison between patients and controls. [1]Subjects with mixed ethnicity and race were counted twice. [2]NIH activity and chronicity indices available in n=124. [3]n=139. [4]Presence of any clinical extrarenal features of the SLE Disease Activity Index (SLEDAI). [5]Predisone dose or equivalence at baseline in n=133, excluding one patient with missing data and seven patients receiving intravenous methylprednisolone ('pulse') dose ranging from 500-1000mg. [6]Includes immunosuppressants and biologics (azathioprine, tacrolimus, cyclosporin, methotrexate, abatacept or belimumab). [7]Renal response was determined at week 52 if baseline UPCR was $\geq$ 1.

**Supplemental Table 2.** Number of samples analyzed with the different panels after filtering for quality control

| | Lupus nephritis patients (Total n = 227) | | | | | Controls (Total n = 40) | | | | |
|---|---|---|---|---|---|---|---|---|---|---|
| **Panels** | **B** | **T** | **M** | **NK** | **4 panels** | **B** | **T** | **M** | **NK** | **4 panels** |
| **Samples** | 224 | 213 | 203 | 191 | 185 | 40 | 40 | 40 | 39 | 39 |
| **Subjects** | 145 | 139 | 134 | 125 | 124 | 40 | 40 | 40 | 39 | 39 |
| **Subjects with baseline visit** | 140 | 131 | 125 | 116 | 115 | 40 | 40 | 40 | 39 | 39 |
| **Subjects with follow-up visits*** | 49 | 46 | 45 | 43 | 42 | 0 | 0 | 0 | 0 | 0 |
| at least bas. and week-12 | 42 | 38 | 35 | 35 | 33 | 0 | 0 | 0 | 0 | 0 |
| at least week-12 and -52 | 34 | 33 | 32 | 30 | 27 | 0 | 0 | 0 | 0 | 0 |
| at least bas. and week-52 | 33 | 31 | 27 | 26 | 23 | 0 | 0 | 0 | 0 | 0 |
| at least bas, week-12 and -52 | 30 | 28 | 25 | 24 | 21 | 0 | 0 | 0 | 0 | 0 |

Samples were stained with panels designed to characterize B cells (B panel), T cells (T panel), myeloid cells (M panel) and NK cells (NK panel). All values represent the number of samples or subjects who had cells stained with each panel or with all 4 panels. If the cell counts in samples were low after thawing, the panels were prioritized as following: B panel first, T panel, M panel and NK panel last. *Include any subjects with any 2 visits (bas. = baseline).

**Supplemental Table 3**. Baseline characteristics and renal response at week 52 of LN patients stratified by immunophenotype subgroups (n=115 LN with samples analyzed with 4 panels)

| | **LN-G0 "control-like"** | **LN-G1 "IFN-I high"** | **LN-G2 "cytotoxic T"** | ***global p val. **** | ***G1-G2 p val. £*** |
|---|---|---|---|---|---|
| **N** | **23** | **46** | **46** | | |
| **Age, median (IQR)** | 41 (31-46) | 30 (24-39) | 33 (26-44) | 0.007 | 0.06 |
| **Female** | 19 (83) | 37 (80) | 42 (91) | 0.31 | |
| **Ethnicity and race** | | | | | |
| **Hispanic or Latino** | 11 (48) | 7 (15) | 19 (41) | 0.006 | 0.01 |
| **Black** | 10/19 (53) | 20/43 (47) | 23/41 (56) | 0.70 | |
| **Previous renal biopsy** | 20 (87) | 34 (74) | 29 (63) | 0.10 | |
| **Proliferative class (III or IV +/- V)** | 9 (39) | 31 (67) | 39 (85) | <0.001 | 0.09 |
| **Activity index[1], median (IQR)** | 1 (0-1) | 4 (0-6) | 5 (3-8) | <0.001 | 0.03 |
| **Chronicity index[1], median (IQR)** | 6 (3-7) | 3 (1-4) | 3 (2-4) | 0.005 | 0.26 |
| **Serum creatinine mg/ml, median (IQR)** | 1.0 (0.8-1.5) | 0.8 (0.7-1.0) | 1.0 (0.8-1.3) | 0.04 | 0.04 |
| **UPCR, median (IQR)** | 3.4 (1.6-4.8) | 1.4 (1.0-2.7) | 2.2 (1.5-4.3) | 0.02 | 0.009 |
| **Presence of cSLEDAI extrarenal[2]** | 5 (26) | 16 (37) | 23 (50) | 0.05 | |
| **Positive anti-dsDNA** | 5 (22) | 38/45 (84) | 36 (78) | <0.001 | 0.63 |
| **Low C3 and/or low C4** | 5 (22) | 31/45 (69) | 34 (74) | <0.001 | 0.62 |
| **Medication** | | | | | |
| **Hydroxychloroquine** | 19 (86) | 42 (91) | 34 (74) | 0.08 | |
| **Prednisone, med (IQR)[3]** | 0 (0-10) | 0 (0-7) | 20 (5-40) | <0.001 | <0.001 |
| **Any immunosuppressants[4]** | 17 (77) | 39 (85) | 30 (65) | 0.08 | |
| **Mycophenolate** | 13 (59) | 32 (70) | 23 (50) | 0.25 | |
| **Renal response at week 52[5]** | | | | | |
| **CR, PR, NR** | 3(18), 3(18), 11(65) | 6(18), 9(27), 19(56) | 15(41), 9(24), 13(35) | 0.04 | 0.03 |

UPCR = urine protein/creatinine ratio. CR, PR, NR = complete, partial and non-responder. Data is presented as N (%) unless specified otherwise. The proportions, median and IQR are calculated on the total number of subject in the group unless specified otherwise. *Chi-square, Fisher exact or Kruskal-Wallis tests when appropriate to test for differences across the three groups. £Chi-square, Fisher exact or Mann-Whitney U test to test for differences between G1 and G2. [1]NIH activity and chronicity indices available in n=100. [2]Presence of any clinical extrarenal features of the SLE Disease Activity Index (SLEDAI). [3]Predisone dose or equivalence at baseline in n=110, excluding five patients receiving intravenous methylprednisolone ('pulse') dose ranging from 500-1000mg. [4]Includes any immunosuppressants and biologics (azathioprine, tacrolimus, cyclosporine, methotrexate, abatacept or belimumab). [5]Renal response was determined at week 52 if baseline UPCR was ≥ 1 (n=88).

**Supplemental Table 4.** Markers, clones, and metals included in each mass cytometry panel

| B panel | | | T panel | | | Myeloid panel | | | NK panel | | |
|---|---|---|---|---|---|---|---|---|---|---|---|
| marker | clone | metal | marker | clone | metal | marker | clone | metal | marker | clone | metal |
| CD45 | HI30 | 89Y | CD45 | HI30 | 89Y | CD45 | HI30 | 89Y | CD45 | HI30 | 89Y |
| CD172ab | SE5A5 | 111Cd | CD172ab | SE5A5 | 111Cd | CD172ab | SE5A5 | 111Cd | CD172ab | SE5A5 | 111Cd |
| CD8a | RPA T8 | 112Cd | CD8a | RPA T8 | 112Cd | CD8a | RPA T8 | 112Cd | CD8a | RPA T8 | 112Cd |
| CD20 | 2H7 | 113Cd | CD20 | 2H7 | 113Cd | CD20 | 2H7 | 113Cd | CD20 | 2H7 | 113Cd |
| CD4 | RPA T4 | 114Cd | CD4 | RPA T4 | 114Cd | CD4 | RPA T4 | 114Cd | CD4 | RPA T4 | 114Cd |
| CD3 | UCHT1 | 115In | CD3 | UCHT1 | 115In | CD3 | UCHT1 | 115In | CD3 | UCHT1 | 115In |
| CD56 | NCAM16.2 | 116Cd | CD56 | NCAM16.2 | 116Cd | CD56 | NCAM16.2 | 116Cd | CD56 | NCAM16.2 | 116Cd |
| CD27 | O323 | 141Pr | CCR6 | G034E3 | 141Pr | Siglec-6 | 767329 | 141Pr | GNLY | Polyclonal | 141Pr |
| Bcl-6 | IG191E/A8 | 142Nd | CD45RA | REA562 | 142Nd | TLR4 | 610015 | 142Nd | KIR2DS1 | 1127B | 142Nd |
| SLAMF7 | 235614 | 143Nd | MX1 | D3W7I | 143Nd | CD36 | 5-271 | 143Nd | CD2 | TS1/8 | 143Nd |
| CD24 | ML5 | 144Nd | CCR4 | L291H4 | 144Nd | CD64 | 10.1 | 144Nd | DAP12 | 406288 | 144Nd |
| CD19 | HIB19 | 145Nd | PU.1 | phpu13 | 145Nd | CD163 | REA812 | 145Nd | NKG2C | REA205 | 145Nd |
| AICDA | EK2-5G9 | 146Nd | SH2D1A | 1A9 | 146Nd | CD74 | LN2 | 146Nd | SH2D1A | 1A9 | 146Nd |
| CD86 | IT2.2 | 147Sm | CD45RO | REA611 | 147Sm | CD86 | IT2.2 | 147Sm | CD7 | 6B7 | 147Sm |
| CD1c | L161 | 148Nd | CXCR3 | REA232 | 148Nd | CD1c | L161 | 148Nd | GZMA | CB9 | 148Nd |
| CD22 | HIB22 | 149Sm | GZMK | GM26E7 | 149Sm | CD1d | 51.1 | 149Sm | GZMK | GM26E7 | 149Sm |
| CD11c | Bu15 | 150Nd | TACTILE | 628211 | 150Nd | CD11c | Bu15 | 150Nd | 2B4 | C1.7 | 150Nd |
| CD5 | UCHT2 | 151Eu | PD-1 | EH12.2H7 | 151Eu | CD123 | 6H6 | 151Eu | TCRVd1 | REA173 | 151Eu |
| Bcl-2 | 100 | 152Sm | CTLA-4 | L3D10 | 152Sm | CD14 | M5E2 | 152Sm | PSGL-1 | CHO131 | 152Sm |
| IgD | IA6-2 | 153Eu | CD69 | FN50 | 153Eu | CD85d | 42D1 | 153Eu | CD69 | FN50 | 153Eu |
| CXCR5 | J252D4 | 154Sm | CXCR5 | J252D4 | 154Sm | CD15 | MC-480 | 154Sm | TCRgd | REA591 | 154Sm |
| CD23 | EBVCS-5 | 155Gd | CD15s | FH6 | 155Gd | Siglec-1 | 7-239 | 155Gd | EOMES | WD1928 | 155Gd |
| CD95 | DX2 | 156Gd | CD8b | SIDI8BEE | 156Gd | XCR1 | 1097A | 156Gd | CD8b | SIDI8BEE | 156Gd |
| CD25 | M-A251 | 157Gd | CD25 | M-A251 | 157Gd | CD16 | 3G8 | 157Gd | CD16 | 3G8 | 157Gd |
| CD39 | A1 | 158Gd | CD39 | A1 | 158Gd | CD39 | A1 | 158Gd | CD39 | A1 | 158Gd |
| TLR9 | S16013D | 159Tb | TCF1 | 7F11A10 | 159Tb | TLR9 | S16013D | 159Tb | PLZF | R17-809 | 159Tb |
| CD307d | 413D12 | 160Gd | ICOS | C398.4A | 160Gd | FPR1 | 350418 | 160Gd | NKp30 | P30-15 | 160Gd |
| CD138 | REA929 | 161Dy | AHR | FF3399 | 161Dy | CD303 | REA693 | 161Dy | MR-1-tet | 5-OP-RU | 161Dy |
| Nur77 | H1648 | 162Dy | Nur77 | H1648 | 162Dy | MARCO | Polyclonal | 162Dy | NKp80 | 5D12 | 162Dy |
| CD83 | HB15e | 163Dy | CCR2 | K036C2 | 163Dy | CCR2 | K036C2 | 163Dy | 4-1BB | REA765 | 163Dy |
| CD79b | CB3-1 | 164Dy | CD161 | HP-3G10 | 164Dy | CD141 | M80 | 164Dy | CXCR6 | K041E5 | 164Dy |
| CD38 | HIT2 | 165Ho | FoxP3 | REA1253 | 165Ho | CD38 | HIT2 | 165Ho | KIR2DS2 | Polyclonal | 165Ho |
| CD40 | 5C3 | 166Er | CD40L | 24-31 | 166Er | CLEC9A | REA976 | 166Er | CD107a | H4A3 | 166Er |
| CD10 | HI10a | 167Er | GZMB | GB11 | 167Er | CD84 | CD84.1.21 | 167Er | GZMB | GB11 | 167Er |
| IgA | IS11-8E10 | 168Er | Helios | REA829 | 168Er | HO-1 | HO-1-1 | 168Er | NKp46 | REA808 | 168Er |
| Pax-5 | 1H9 | 169Tm | CX3CR1 | REA385 | 169Tm | CX3CR1 | REA385 | 169Tm | CD3z | 6B10.2 | 169Tm |
| PD-L1 | 29E.2A3 | 170Er | RORyt | REA278 | 170Er | PD-L1 | 29E.2A3 | 170Er | iNKT | 6B11 | 170Er |
| IgG | G18-145 | 171Yb | CD127 | eBioRDR5 | 171Yb | CD206 | 19.2 | 171Yb | TCRVd2 | REA771 | 171Yb |
| ISG15 | 539442 | 172Yb | GATA3 | REA174 | 172Yb | IRF8 | REA516 | 172Yb | NKG2D | 149810 | 172Yb |
| CD21 | Bu32 | 173Yb | TIGIT | MBSA43 | 173Yb | CD170 | 1A5 | 173Yb | TCRab | T10B9.1A-31 | 173Yb |
| Ki67 | 8D5 | 174Yb | Ki67 | 8D5 | 174Yb | Ki67 | 8D5 | 174Yb | Ki67 | 8D5 | 174Yb |
| T-bet | 4B10 | 175Lu | T-bet | 4B10 | 175Lu | CD85i | 586326 | 175Lu | Tbet | 4B10 | 175Lu |
| IgM | MHM-88 | 176Yb | CCR7 | REA546 | 176Yb | CD68 | Y1/82A | 176Yb | Perforin | dG9 | 176Yb |
| Blimp1 | 646702 | 194Pt | CD57 | REA769 | 194Pt | CD180 | MHR73-11 | 194Pt | CD57 | REA769 | 194Pt |
| HLA-DR | L243 | 195Pt | HLA-DR | L243 | 195Pt | HLA-DR | L243 | 195Pt | HLA-DR | L243 | 195Pt |
| CD52 | HI186 | 196Pt | CD103 | Ber-ACT8 | 196Pt | FOLR2 | 94b/FOLR2 | 196Pt | SLAMF6 | 292811 | 196Pt |
| IgE | MHE-18 | 198Pt | CD38 | HIT2 | 198Pt | CD115 | 61708 | 198Pt | CD38 | HIT2 | 198Pt |
| CD11b | ICRF44 | 209Bi | CD11b | ICRF44 | 209Bi | CD11b | ICRF44 | 209Bi | CD11b | ICRF44 | 209Bi |
