## Supplemental Figures for "Blood immunophenotyping identifies distinct kidney histopathology and outcomes in patients with lupus nephritis"

### Supplemental Figure 1

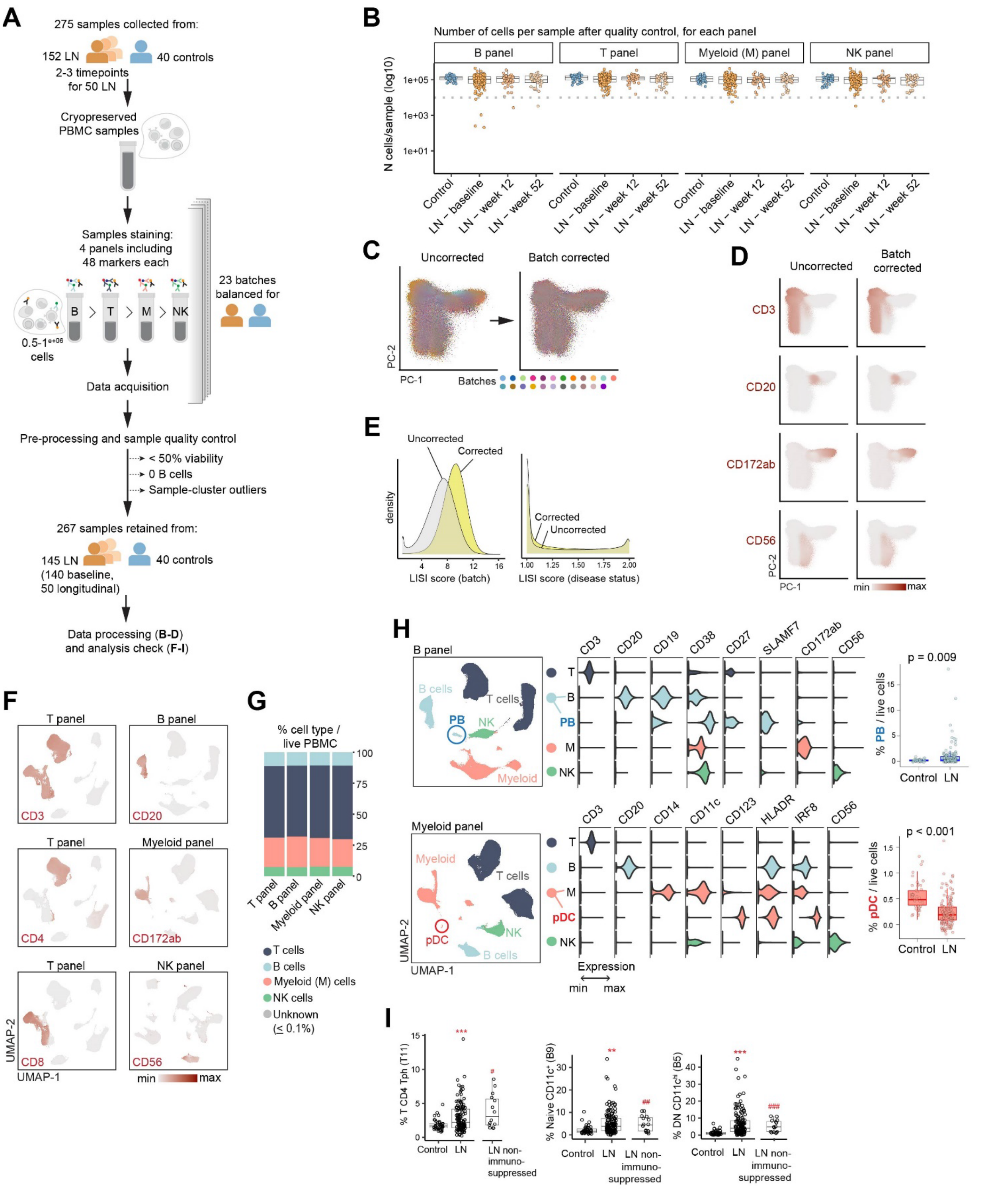

**Supplemental Figure 1. Overview of blood immunophenotyping by mass cytometry in a cohort of lupus nephritis (LN).** (**A**) Sample processing and quality control pipeline. (**B**) Number of cells analyzable, after filtering for quality control, in each panel and stratified by disease status and timepoints. The dotted grey line represents the threshold of 10,000, used for downsampling. (**C**) Representative example of distribution of cells stained with the T panel in the first two principal components, before and after batch correction. (**D**) Expression of key markers before and after batch correction in the T panel. (**E**) Local inverse Simpson's index (LISI) scores per cell in the T panel measuring the mixture of cells per batch or disease status (SLE versus control). Increased index represents increased mixture of cells. (**F**) UMAP representing the expression of key lineage markers by cells stained in the four different panels. (**G**) Proportion of the main cell types, amongst total peripheral blood mononuclear cells, identified in each panel. (**H**) Identification of plasmablasts and plasmacytoid dendritic cells in the UMAP space and by marker expression, with expected proportions when comparing all baseline LN patients (n=140 in the B panel and n=125 in the myeloid panel) and controls (n=40) (Wilcoxon signed-rank test). (**I**) Comparison of proportion of specific cell subsets (% Tph amongst T cells and % of $CD11c^{+}$ B cells amongst B cells) between 40 controls and total LN patients (**p<0.01, ***p<0.001 by linear regression adjusting for age, sex, ethnicity and race) or between controls and 15 LN without immunosuppression and prednisone $\leq$ 5mg ($^{\#}$p<0.05, $^{\#\#}$p<0.01, $^{\#\#\#}$p<0.001 by Wilcoxon signed-rank test).

#### Supplemental Figure 2

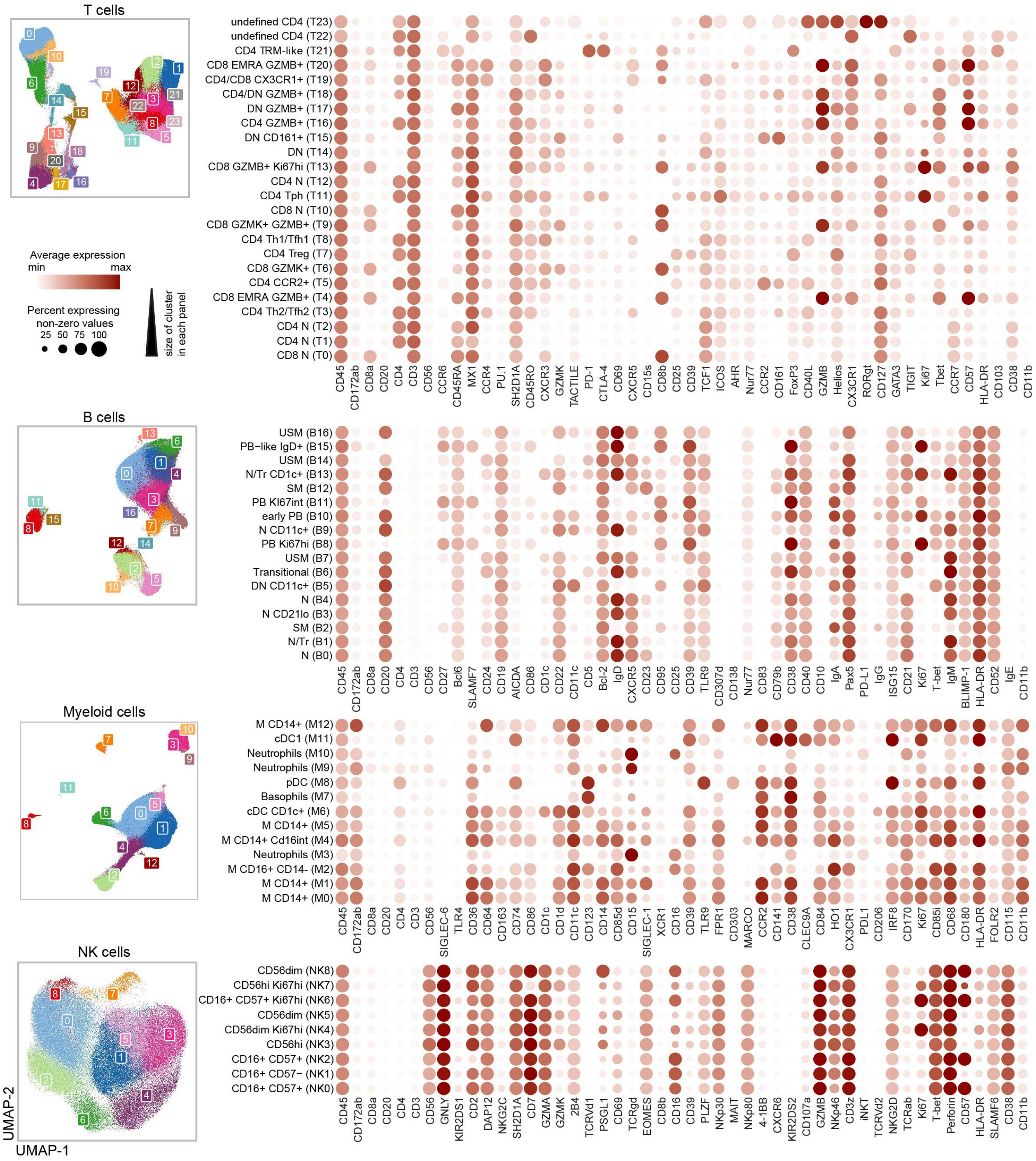


**Supplemental Figure 2. Cell-type specific clustering and marker expression per cluster.** Cell type specific clusters colored in the UMAP space. All proteins included in each panel and used for clustering cells are shown in the dotplots. Clusters are ordered by size (number of cells) from bottom to top, for each panel.

#### Supplemental Figure 3

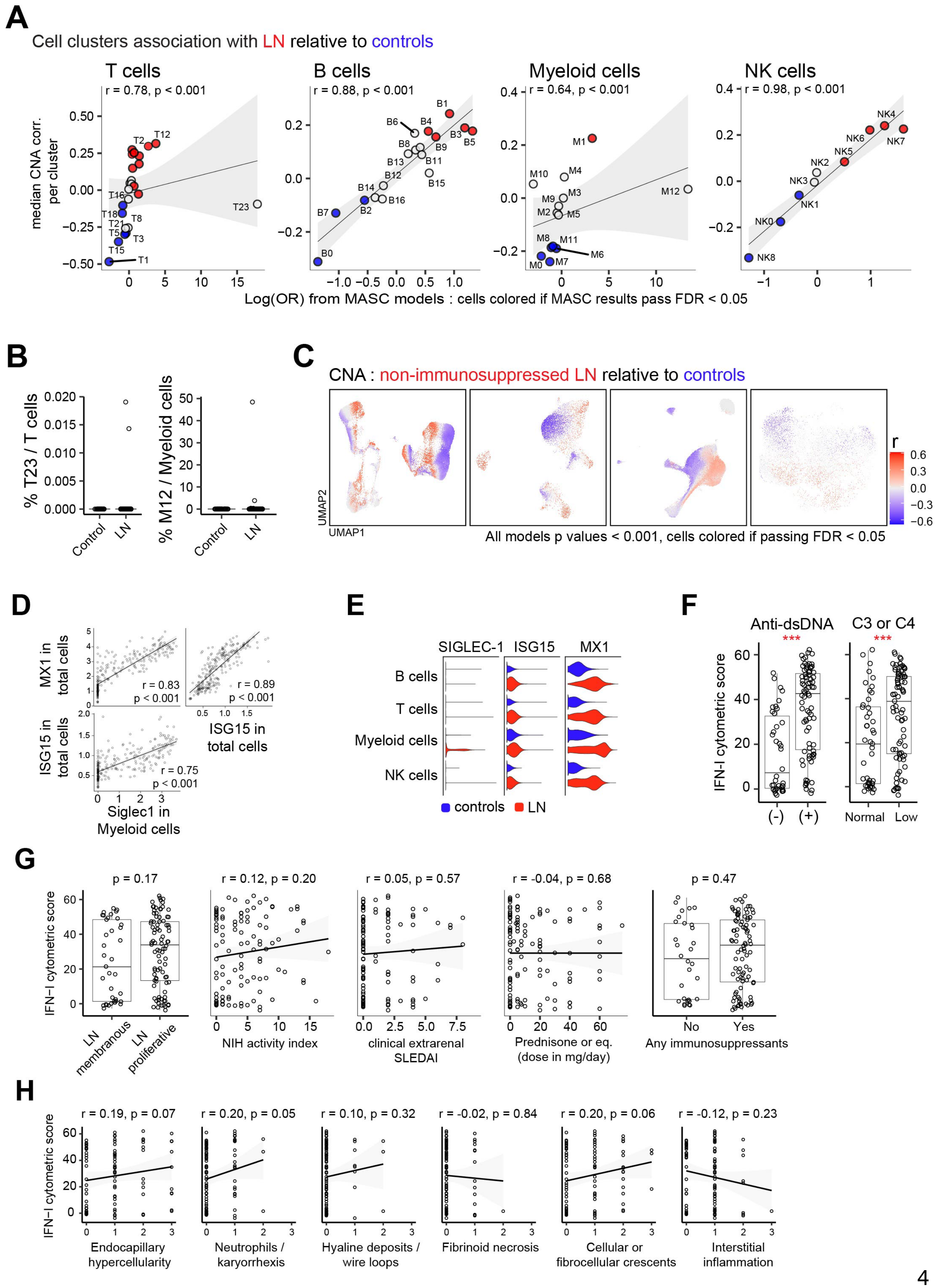

**Supplemental Figure 3. Comprehensive identification of immune alterations in lupus nephritis (LN) patients reveal a proteomic type I interferon signature**. (**A**) Identification of cell clusters associated with LN (max n=140 in the B panel and min 116 in the NK panel) relative to controls (max n=40 in all panels except for n=39 in the NK panel), using two different approaches : covarying neighborhood analysis (CNA) and a single-cell mixed-effect model (MASC). Plots represent the correlation between the two models using Spearman's rho. (**B**) Example of two cellular cluster outliers identified as outliers in figure a. (**C**) Identification of cells associated with a subgroup of LN patients without immunosuppressive therapy and prednisone dose maximum 5 mg at the time of sample collection (max n=15 in the B panel and min n=13 in the NK panel), relative to controls. (**D**) Correlation between the median expression per sample of MX1 and ISG amongst total live cells and SIGLEC-1 amongst myeloid cells (Spearman's rho correlation). (**E**) Distribution of the level of expression of type I interferon induced proteins in major cell types. (**F**) Comparison of a combined type I interferon score (sum of normalized MX1 and ISG15 in live cells and SIGLEC-1 in myeloid cells) in LN patients with serologic parameters. (**G**) Lack of association between cytometric type I interferon score and clinical or histologic characteristics, (**H**) including with the NIH activity subscores, using Wilcoxon rank-sum test or Spearman's rho correlation.

**Supplemental Figure 4**

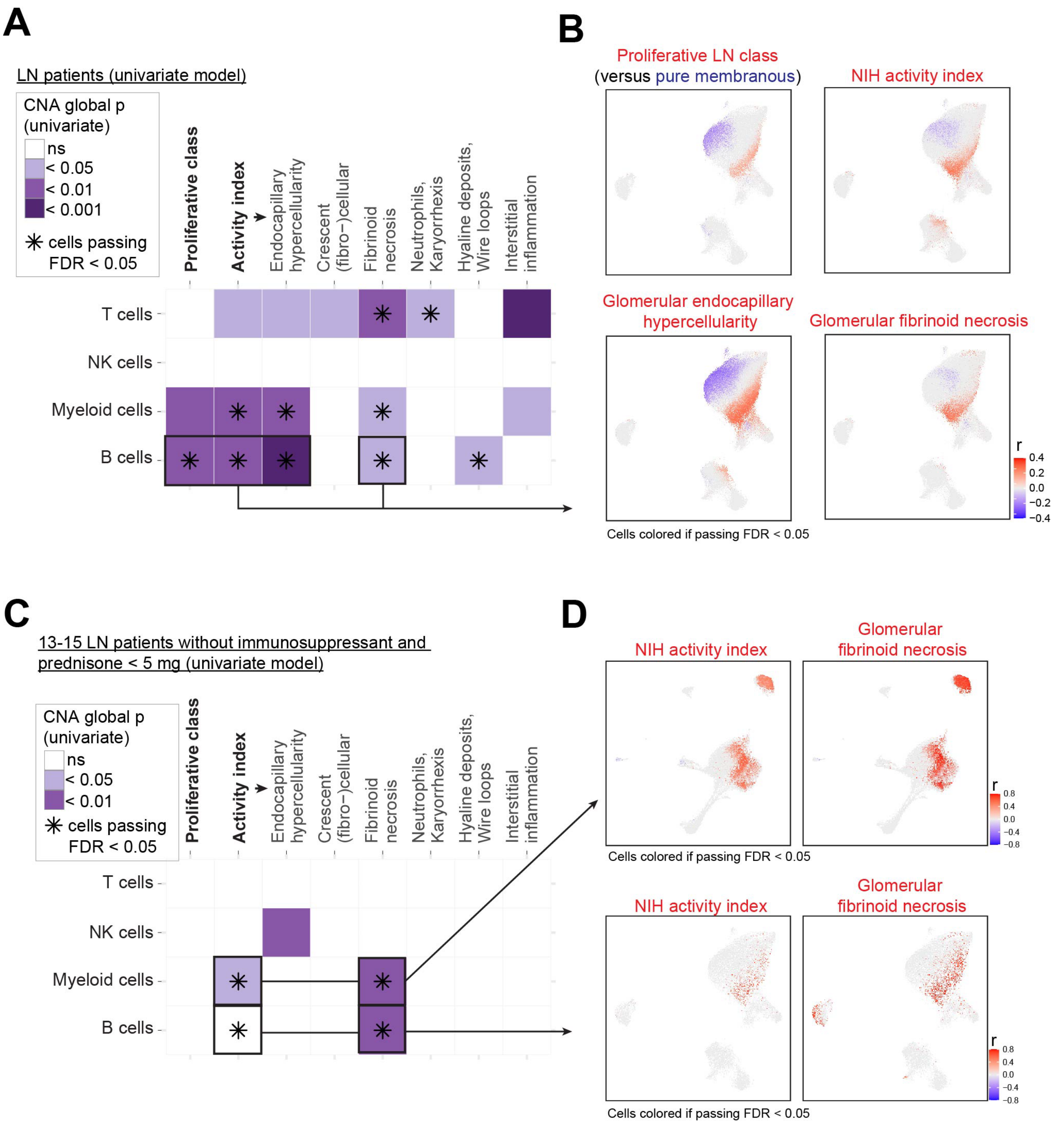


**Supplemental Figure 4. Circulating cell-type specific alterations are associated with histologic patterns of active LN.** (**A**) Summary of the results testing the association within each blood cell type (y axis) and different histologic patterns of LN disease (x axis). Statistical significance mentioned in the graph were determined using a univariate covarying neighborhood analysis (CNA). (**B**) Selection of detailed CNA results between B cell alterations and specified histologic characteristics. (**C**) Summary of CNA results including only LN patients with no immunosuppressive therapy and prednisone $\leq$ 5mg with (**D**), representative detailed results in the myeloid and B panels.

#### Supplemental Figure 5

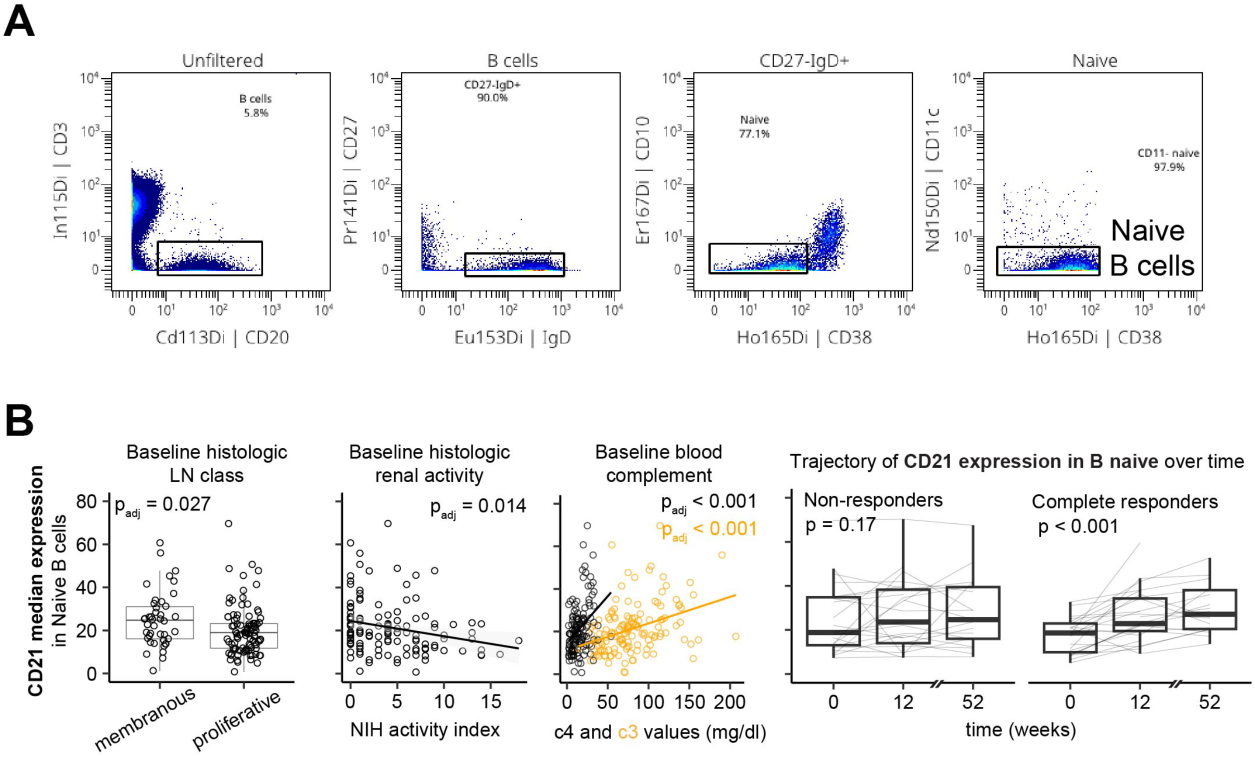


**Supplemental Figure 5. Naïve B cells shift their phenotype towards a low expressing CD19 and CD21 profile in active and proliferative LN patients.** (**A**) Gating strategy to identify naïve B cells by mass cytometry. (**B**) Median CD21 expression amongst naïve B cells association with histological characteristics, complement values (C3 and C4) and longitudinal change over time. Statistical significances were determined either using Wilcoxon sum-rank test (LN class), Spearman's rho correlation (NIH activity index and complement) or a mixed effect model with patient as a random effect (changes over time).

#### Supplemental Figure 6

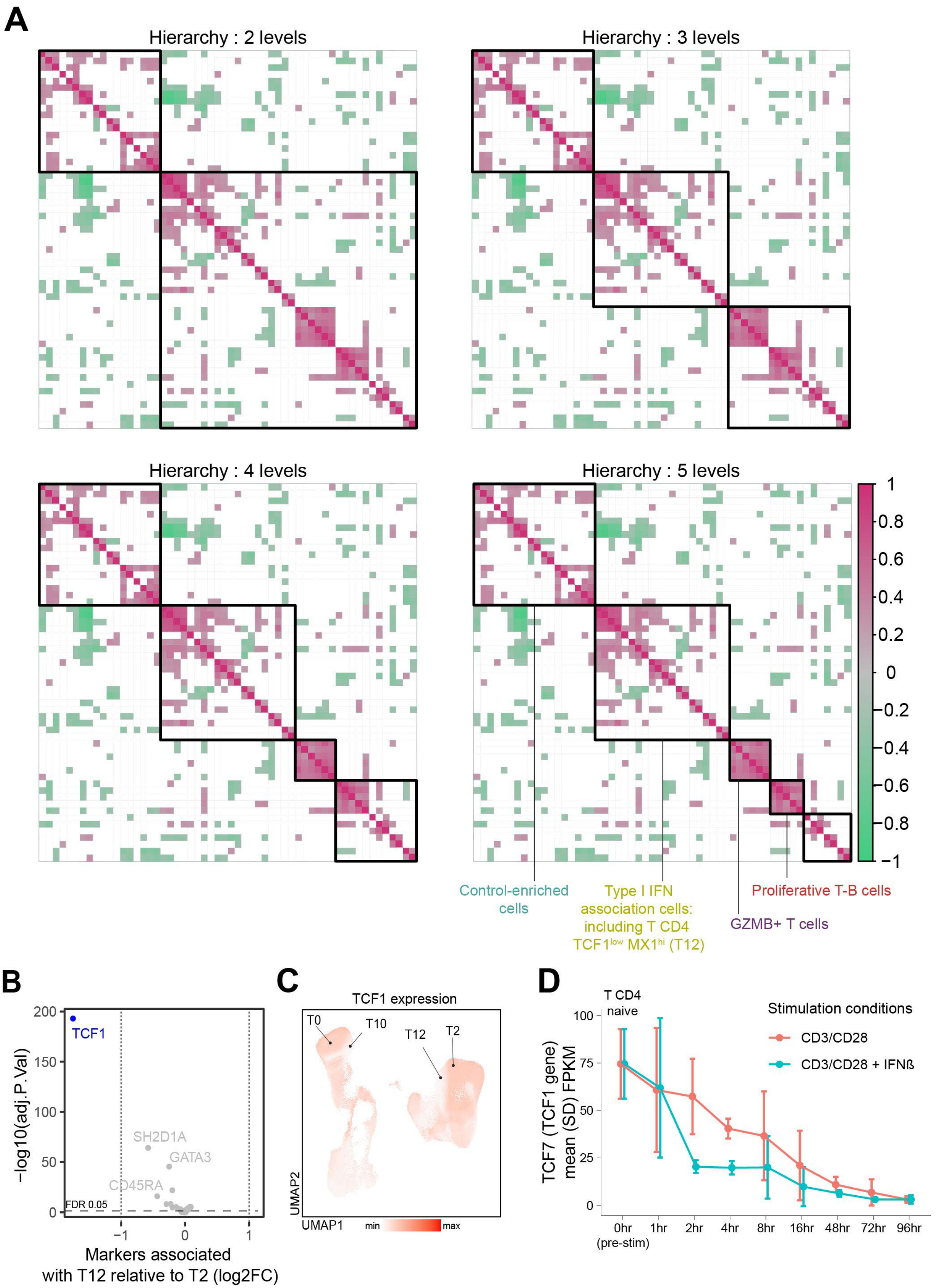

**Supplemental Figure 6. Hierarchical clustering of co-correlated cells identifies cell subsets strongly associated with type I interferon signaling. (A**) Correlation between 55 immune cell subsets organized by hierarchical clustering algorithm. The black boxes represents the first 5 levels of hierarchy. Main co-correlating sets of cells are labeled by the main characteristics of cells including in the groups. (**B**) Differential expression of markers included in the T panel between the cell subset T2 and T12. Statistical significance was determined using the limma package. (**C**) UMAP of TCF1 expression in T cells. (**D**) Changes in TCF7 (TCF1 gene) expression in healthy naïve CD4 T cells stimulated with or without interferon-$\beta$, using published bulk RNA-seq dataset.

### Supplemental Figure 7

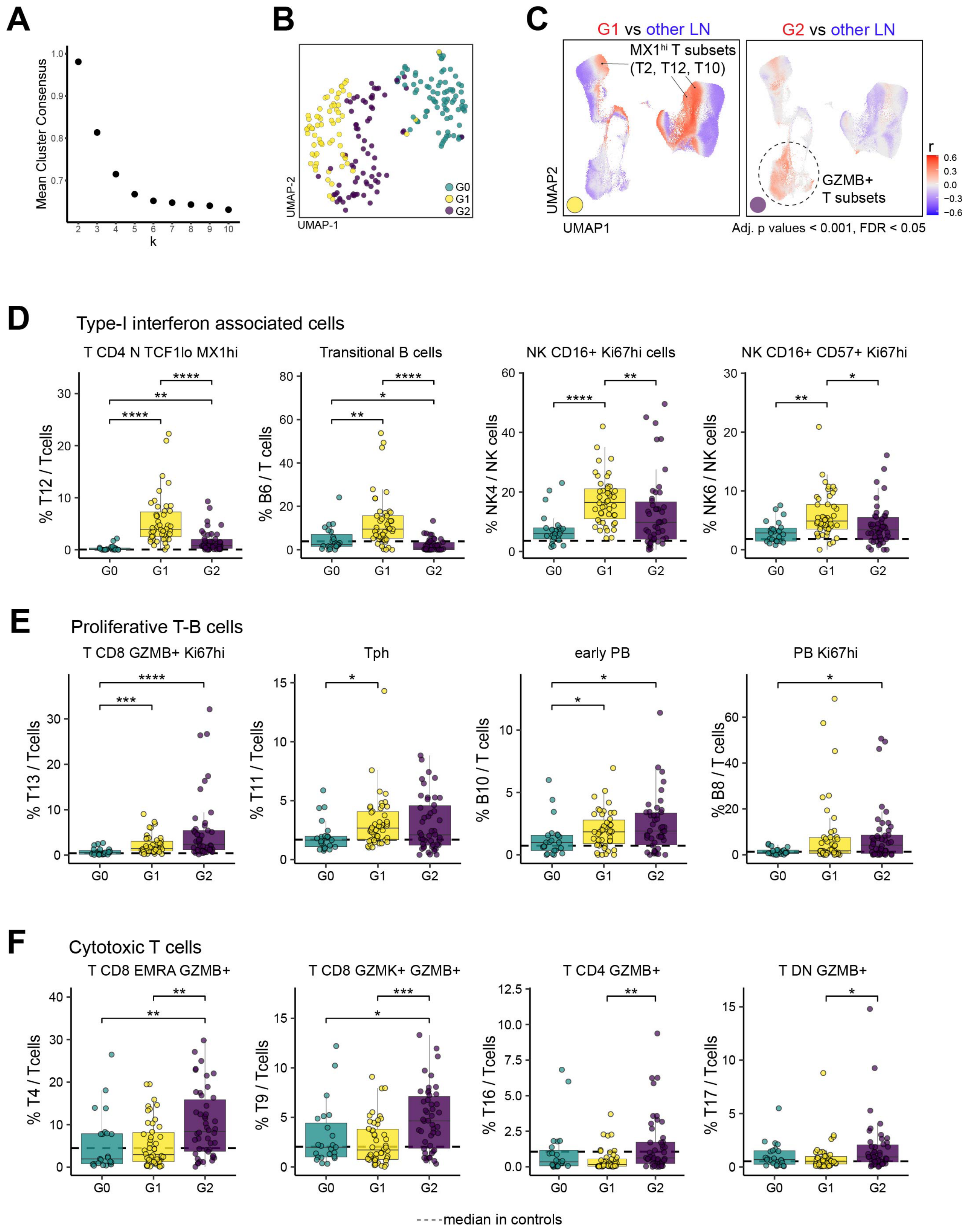

**Supplemental Figure 7. Circulating immune cell subsets characterizing three LN groups. (A)** Stability of sample membership to clusters depending on the number of clusters by repeating 1000 K-means clustering on a resampled dataset without replacement. (**B**) UMAP distribution of all samples included in this study (n=267) based on the proportion of 55 immune cell subsets. (**C**) Cell neighborhood associations with the K-means defined groups G1 and G2 relative to the other groups. (**D-F**) Comparison of selected cell subsets between the three K-means defined groups of LN patients at baseline (23 G0, 46 G1 and 46 G2). Statistical significance was determined by Kruskal-Wallis test followed by Dunn's multiple comparisons.

**Supplemental Figure 8.**

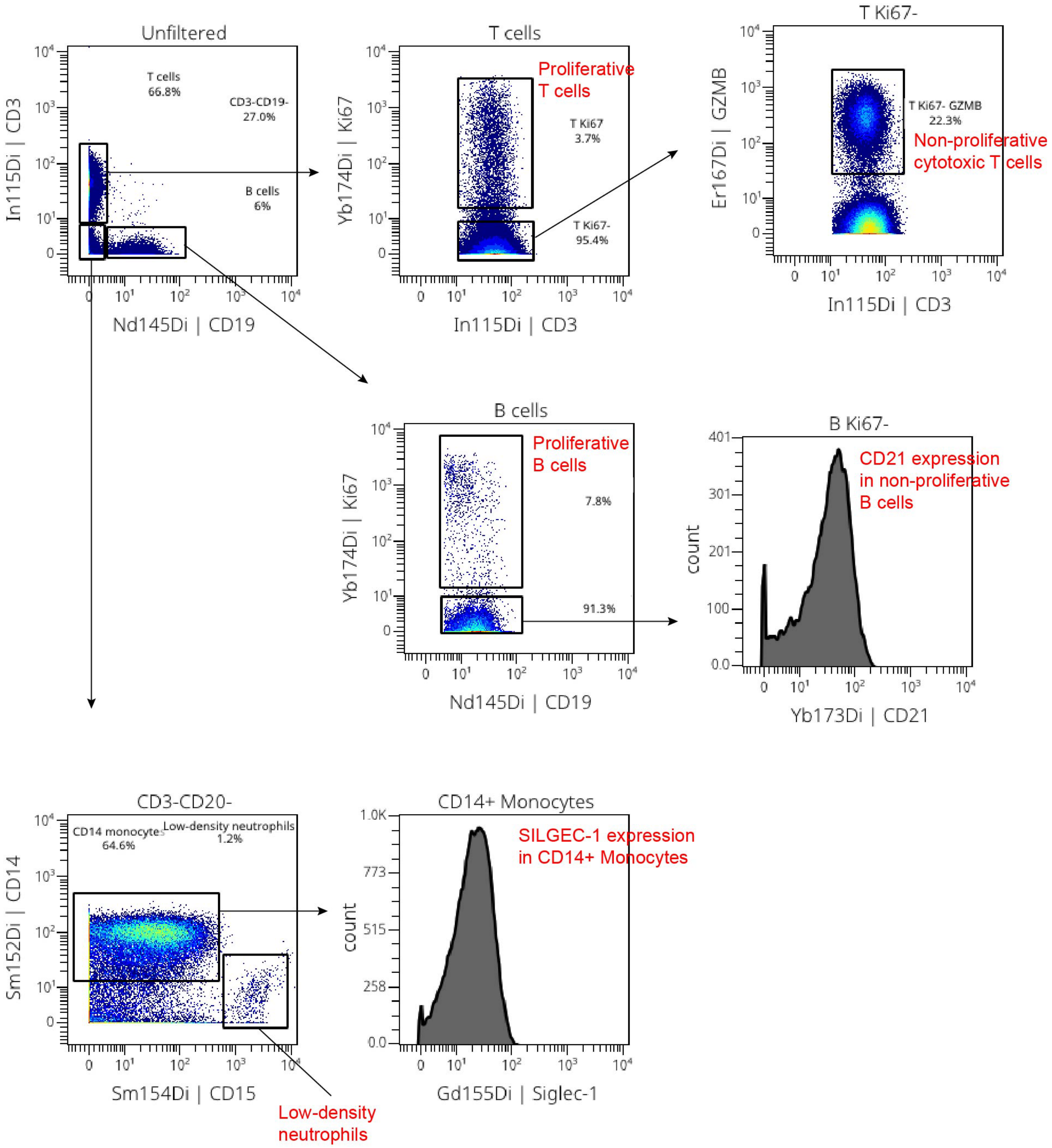


**Supplemental Figure 8. Manual gating strategy to obtain simplified cellular immunophenotype.**
