## Supplemental Methods for "Blood immunophenotyping identifies distinct kidney histopathology and outcomes in patients with lupus nephritis"

### Detailed graph-based clustering, visualization of protein expression and cell subset annotation

After batch correction, we built a nearest-neighbor graph using the function FindNeighbors using the 20 first harmonized PCs. The clustering was performed using a Louvain-based algorithm implemented in the FindCluster function (Seurat package). For this step we used a resolution of 0.5 for all panels after confirming that all major cell subsets could be identified. We projected cells into two dimensions using the runUMAP function based on the 20 first harmonized PCs, and with the following arguments: metric = cosine, min.dist = 0.08, spread = 1. We obtained between 23 to 29 clusters in each panel. We annotated each cluster based on common lineage markers and additional markers specific to each panel as either: B cells (including clusters with a plasmablast phenotype), T cells, myeloid/DC cells (including clusters with a plasmacytoid dendritic phenotype, basophil phenotype, and neutrophil phenotype) and NK cells. Clusters that were not attributed clearly to one of these cell types were labeled undetermined and were excluded from further analysis (< 0.1% of total live cells for each panel) (**Supplemental 1F-H**).

As a second step, we extracted the cell type of interest in the dedicated panel (e.g., T cells in the T panel) and re-clustered the data. We applied a similar pipeline for each subsetted Seurat object by applying again the functions: RunPCA(), RunHarmony(), FindNeighbors(), FindCluster() and RunUMAP(). Additionally, for the cell-type specific nearest-neighbor graph, we used a k=30 based on an elbow plot of the within-cluster sum of squares. For the Louvain-based algorithm, we optimized resolution for each cell type (0.8 for B cells, 0.7 for T cells, 0.3 for Myeloid cells, 0.3 for NK cells), based on the cluster distribution and manual check of expression of critical proteins in each cluster to gain the biological interpretations that made the most sense. Cell-type specific clusters were labeled with a first letter corresponding to the panel they were extracted from, and a number based on the cluster size (e.g., the cluster “B0” corresponds to the largest cluster within

B cells extracted from the B panel) and were further annotated based on canonical markers expression and relevant literature.

Overall, for visualization of protein expressions, we showed the arcsinh-transformed data in multiple plot types adapted from the Seurat package: 1) Dotplots to display all proteins in each cluster with dots colored by average of unscaled expression and sized by percentage of cells with non-zero values; 2) Violin plots to show the distribution of a selection of proteins, where the minimal and maximal thresholds were data-driven by the cells expressing the lowest and highest expression; 3) Feature plots to map a selection of protein expression in the UMAP space, with the same min-max threshold definition than in 2).

**Detailed association testing of cell subsets with disease status and characteristics**

To examine cell subsets associated with LN disease or with clinical conditions within patients with LN, we applied co-varying neighborhoods analysis (CNA)(41) to our dataset using its r version (rcna package, version 0.0.99) implemented in the Seurat package through the association.Seurat function. CNA defines data-driven neighborhoods (here referring to small regions of cells sharing proteomic similarities) and measures the relative abundance of cells from each sample across all neighborhoods. By further applying PCA to the neighborhood abundance matrix, CNA identifies dominant axes of variation (NAM-PCs) in cell neighborhood abundance across samples and further allows to test for clinical association including potential covariates, in a linear model. We used this approach to test cell-association with individual clinical characteristics in a univariate way and by adjusting for potential confounding factors, as specified in the results and legends. For each model, we examined and reported the global CNA p-value, which was defined through a permutation test, and further evaluated the cell-neighborhoods that passed a threshold of FDR<0.05. To visualize the local associations with the tested variables, we mapped the results in the UMAP space and colored the cell neighborhoods in red or blue, for any significant positive or negative correlation, respectively, passing the FDR threshold.

We tested the consistency of our results by using a separate single-cell cluster-based approach based on a mixed-effect model (MASC)(42). The model evaluates whether the tested variable (LN relative to controls) improves the null model determining the membership of each cell to its cluster while controlling for inter-individual variability (random effect) and covariates (fixed effects). To run MASC function, we extracted the components from the Seurat object to build a dataframe with one row per cell and the random and fixed variables as columns. MASC results provided the significance and effect size (defined by an odds ratio) of disease status (LN relative to controls) for each cluster. To examine the consistency between the overall results of the 2 approaches, we tested the correlation between the median cell correlation coefficient, defined by CNA, for each cluster, and the odds ratio defined by MASC.

**Detailed identification of circulating immune cell signatures**

We extracted the proportions of each cell subsets among cell type for each sample. To prevent signals from small cluster outliers, we removed cell subset clusters that included more than 50% of samples with a zero value (examples shown in **Supplemental Figure 3B**). We excluded three T cell clusters (T21, T22, and T23), three B cell clusters (B14, B15, and B16) and two myeloid clusters (M10 and M12). We obtained a matrix of 55 cell subset (=cell clusters) proportions with sample representing the rows and columns representing cell subsets. We studied the correlation between all cell clusters and the type I interferon score, including all baseline samples from patients with LN stained with four panels (n=115). To obtain the Spearman's rho correlations coefficients and p values between all cell subsets and type I interferon score, we used the rcorr function from the Hmisc package (version 5.1-1). We applied FDR correction to the p values to account for multiple testing using the p.adjust function from the stats package (version 4.3.1). We then used the R package corrplot (version 0.92) to explore visually our matrix of correlations by highlighting the correlations that passed FDR and by reordering the correlation by hierarchical clustering using the Ward distance method (**Figure 3A, Supplemental Figure 6A**).

### Detailed study approval

All participants provided written informed consent before study enrollment, and human study protocols were approved in accordance with the Declaration of Helsinki by the institutional review boards (IRBs) at each participating sites, which included: Johns Hopkins University, New York University, University of Rochester Medical Center, Oklahoma Medical Research Foundation, University of Cincinnati, Albert Einstein College of Medicine, University of California San Francisco, Northwell Health, Medical University of South Carolina, Texas University El Paso, University of Michigan, University of California San Diego, University of California Los Angeles, Cedars-Sinai Medical Center.
